# Mutational and functional heterogeneity of homology-directed repair deficiency and clinical implications

**DOI:** 10.64898/2026.05.27.725139

**Authors:** Salome J Zhao, Gene C C Koh, Rachel Brough, Dragomir B Krastev, Veenu Tripathi, Feifei Song, Sheyne Choi, Daniella Black, Soraya Boushaki, Cherif Badja, Rebecca A Dagg, Giuseppe Rinaldi, Ping Jing Toong, Andrea Degasperi, Rohit Prakash, Scott Shooter, Jan Czarnecki, Yasin Memari, Helen R Davies, Stephen J Pettitt, Madalena Tarsounas, Maria Jasin, Andrew Tutt, André Nussenzweig, Christopher J Lord, Serena Nik-Zainal

**Affiliations:** Department of Genomic Medicine, School of Clinical Medicine, University of Cambridge, Cambridge CB2 0QQ, UK; Sir Jeffrey Cheah Sunway Medical School, Faculty of Medical and Life Sciences, Sunway University, Sunway City, Malaysia; Precision Oncology Laboratory, The Breast Cancer Now Toby Robins Research Centre, The Institute of Cancer Research, London SW3 6JB, UK; Laboratory of Genome Integrity, National Cancer Institute, NIH, Bethesda, MD, USA; Genome Stability and Tumorigenesis Group, Department of Oncology, University of Oxford, Oxford OX3 7DQ, UK; Memorial Sloan Kettering Cancer Center, New York, NY 10065, USA

**Keywords:** Mutational signatures, CRISPR screen, homologous recombination repair, synthetic lethality, isogenic cell lines, PARP inhibitor resistance, therapeutic vulnerability, BRCA, DNA repair, mutagenesis

## Abstract

Homology-directed repair deficiency (HRd) encompasses mutations in multiple genes yet is treated clinically as a single entity. Here, through parallel analyses of isogenic knockouts of multiple HR pathway genes, integrating multi-omic analyses with genome-wide CRISPR-Cas9-dependency and resistance screens, we show that HRd is not a single entity but exists along a molecular and functional continuum. *BRCA1, BRCA2, PALB2, RAD51C,* and *RAD51D* mutants shared many HRd-associated mutational signatures, while *RAD51B, BRIP1*, *CDK12* exhibited distinct genomic patterns. Functional heterogeneity was equally apparent: synthetic lethal interactions including *CIP2A* and a novel dependency on *PRDX1* were penetrant across most HRd genotypes, whereas *FANCM* dependency was linked to HRd subtypes characterized by tandem duplications. PARPi resistance screens in distinct HRd contexts uncovered *BRIP1* and *RECQL5* as new *BRCA2*-specific resistance genes. HRd is thus a complex continuum, underscoring why modernizing the molecular taxonomy utilizing all genomic features available per patient is crucial to informing precision interventions.

## INTRODUCTION

Inherited mutations in *BRCA1*/*BRCA2* are associated with widespread genomic instability in tumors, primarily due to defects in homology-directed repair (HR)^1^. Earliest reports identified several phenotypic features including copy number aberrations or “genomic scars” termed “*BRCAness*”^2,3^ in tumors from germline mutation carriers, which were sometimes observed in sporadic cancers. Later, the advent of cancer whole-genome sequencing (WGS) identified high-resolution patterns of mutagenesis or *mutational signatures* in tumors from patients with inherited *BRCA1/BRCA2* mutations that were likewise observed in sporadic cancers. The mutational signatures associated with *BRCA1/BRCA2*-mutant tumors include signatures of single-base substitutions (SBS3 and SBS8), indel signatures characterized by microhomology (InD6), structural variation (SV) signatures (R3 and R5), and widespread copy number alterations, including whole-arm and whole-chromosome loss of heterozygosity (LOH)^1,4–6^. Subsequently, tumors with biallelic loss of *BRCA1*/*BRCA2*, irrespective of the mechanisms of inactivation – whether germline, somatic and/or epigenetic (*e.g., BRCA*1 promoter silencing) *–* were also shown to carry identical mutational signatures. Additionally, biallelic loss of *PALB2* and promoter hypermethylation of *RAD51C* in human cancers were found to manifest these signatures^7^.

In clinical settings, *BRCAness* scars underpin the FDA-approved companion diagnostic assay Myriad MyChoice HRD, used as a biomarker of HR-deficiency (HRd) and for the identification of patients predicted to benefit from selected therapeutics such as platinum salts (Pt-salts) or synthetic lethal compounds such as poly(ADP-ribose) polymerase inhibitors (PARPi)^2,8,9^. Despite these advances, clinical responses are heterogeneous^10–12^: some patients with apparent mutations in *BRCA1/BRCA2* and/or HRd genomic scars appear *de novo* resistant to Pt-salts or PARPi, whereas others that lack classical *BRCAness* exhibit sensitivity to PARPi. What remains unclear is whether all forms of HRd are molecularly or functionally equivalent and whether variation in HRd could at least partially explain the variable efficacy observed with targeted agents. For example, mutations in many HR-related genes are used to select prostate cancer patients for olaparib treatment, yet the relative clinical benefit in *BRIP1*-mutant patients compared to *BRCA1/2*-mutant patients is variable^13,14^. Resistance mechanisms across *BRCAness* cases may also differ by genotype. For example, while *BRCA1*-mutant cancers can develop resistance to PARPi through the loss of 53BP1/Shieldin complex^15,16^, this is not a recognized resistance mechanism seen in *BRCA2*-mutant cancers^17^. The full landscape of genotype-specific resistance routes remains poorly characterized^18^. Taken together, these observations suggest that molecular and functional heterogeneity of HRd has direct therapeutic consequences.

The hypothesis that HRd is not a single entity has remained largely unexplored^19^. A major obstacle to studying this possibility has been the lack of isogenic experimental systems that enable direct, systematic comparison of distinct HRd drivers within a singular, controlled genetic background. Furthermore, although many HR-associated genes are recurrently mutated in patient tumors and HRd-related mutational signatures are relatively prevalent in human tumors, naturally occurring cancer cell lines harboring deleterious *loss-of-function* mutations in HR genes beyond *BRCA1/BRCA2* are rare (*e.g., PALB2, RAD51* paralogs), precluding systematic interrogation of their contribution to HRd. Here, we hypothesize that there is diversity in mutational and functional phenotypes associated with different genes in the same pathway. To assess this hypothesis, we created an isogenic cellular resource of CRISPR-edited genes in the HR pathway and used this series to comparatively evaluate mutational diversity through mutation accumulation experiments, and functional diversity through genome-wide CRISPR-Cas9 *loss-of-function* screens. Below, we dissect several observations and present conceptual advances, with direct implications for biomarker design, therapeutic targeting, and patient stratification.

## RESULTS

### Spectrum of mutational signatures of genes involved in HR

We generated a series of CRISPR-Cas9 knockouts (KOs) targeting canonical HR pathway genes (*BRCA1, BRCA2, PALB2*), RAD51 paralogs (*RAD51B, RAD51C, RAD51D*), and putative HR genes *BRIP1*^20^ and *CDK12*^21,22^ on a common genetic background of *TP53-null* hTERT-immortalized RPE1 cells (**Figure 1A**). We verified the loss of protein expression via immunoblotting, the functional status of the KO models using a plasmid HR reporter assay^23^, and drug sensitivities to Pt-salt cisplatin (**Figures S1A, S1B, and S1C**). High-depth WGS following a 45-day mutation accumulation experiment demonstrated that all KO subclones, except for *ΔBRIP1*, exhibited elevated substitution, indel, and rearrangement counts compared to an unedited control (WT) reporting background mutagenesis. Rearrangement counts were modest overall (**Figure 1B**). To assess qualitative differences, we compared the mutational profiles of KO subclones to bootstrapped WT profiles across a range of mutational loads. Focusing on *ΔBRCA1, ΔBRCA2, ΔPALB2,* and *ΔRAD51C* as they represent recognized HRd states in human cancers^5,7^, subclones of these genotypes exhibited substitution and indel profiles clearly discernible from the unedited WT (**Figure 1C**). We subtracted the background profile from each KO genotype to reveal genotype-specific HRd substitution and indel mutational patterns, which were highly consistent with each other (cosine similarity>0.90) (**Figure 1D; Table S1**). We thus averaged them to derive “ground truth” experimental HRd signatures of substitutions (eSBS.HRd) and indels (eInD.HRd) (**Figure 2A; Tables S2 and S3**), which closely matched cancer-derived SBS3 and InD6^6^ (cosine similarity: 0.90 and 0.99, respectively). Low rearrangement counts precluded conventional 32-channel rearrangement signature analysis. However, collapsing the rearrangement profiles into 16 channels revealed gene-specific differences: *ΔBRCA2* exhibited deletion-dominated profiles (1–10 kb), while *ΔBRCA1* displayed a mix of deletions and short tandem duplications (TD) (1–10 kb) – mirroring R5 deletion and R3 TD rearrangement signatures in HRd tumors^1^, respectively (**Figure 1D**). To reinforce our observations, we analyzed independent subclones of *BRCA1* KO in HAP1 (a cancer cell line) and *BRCA2* KO in MCF10A^24^ (a non-tumorigenic breast epithelial cell line). Both showed eSBS.HRd, eInD.HRd, and gene-specific rearrangement patterns (*i.e.,* TD-dominance in *ΔBRCA1* and deletion-dominance in *ΔBRCA2*), confirming the robustness and universality of HRd signatures, recapitulated in alternative normal and cancer cell models (**Figures S3A and S3B; Tables S4, S5, and S6**).

**Figure 1.**
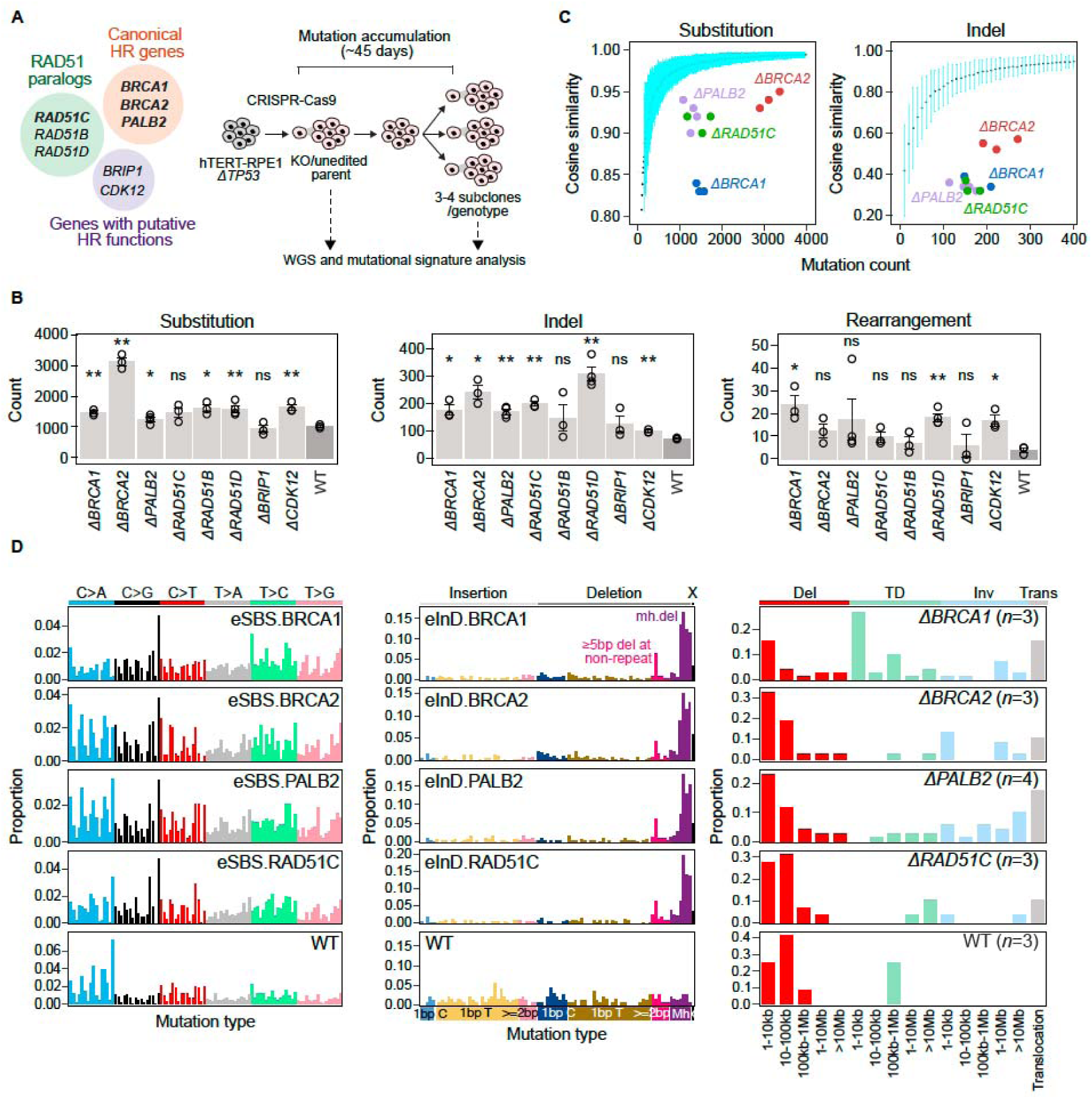
Isogenic HR gene knockouts reveal genotype-specific mutational signatures (A) Schematic of mutation accumulation experiment in *TP53*-null hTERT-immortalized retinal pigment epithelial cell (hTERT-RPE1*^TP^*^53-null^, herewith referred to as WT) with CRISPR knockouts of HR genes. (B) *De novo* mutation burdens of knockouts (*n*=3-4 subclones per genotype; **Tables S2, S3, and S6**). Error bars represent mean□±□SEM. Two-tailed Welch’s t test versus WT; *p*≤0.05 (*), *p*≤0.01 (**). (C) Distinguishing mutational profiles of edited subclones from WT background. Cyan error bars depict mean□± 3□SEM of cosine similarities between *n*□=□100 bootstrapped mutational profiles of unedited WT and the background profile aggregated from *n*□=□7 unedited subclones. The *x*-axis shows the respective mutation count. (D) Background-subtracted mutational signatures of HR gene knockouts. Aggregated mutation profiles are shown for rearrangements due to low burden; *n* denotes the number of subclones per genotype. See also Figures S1 and S2; Tables S2, S3, and S6.

**Figure 2.**
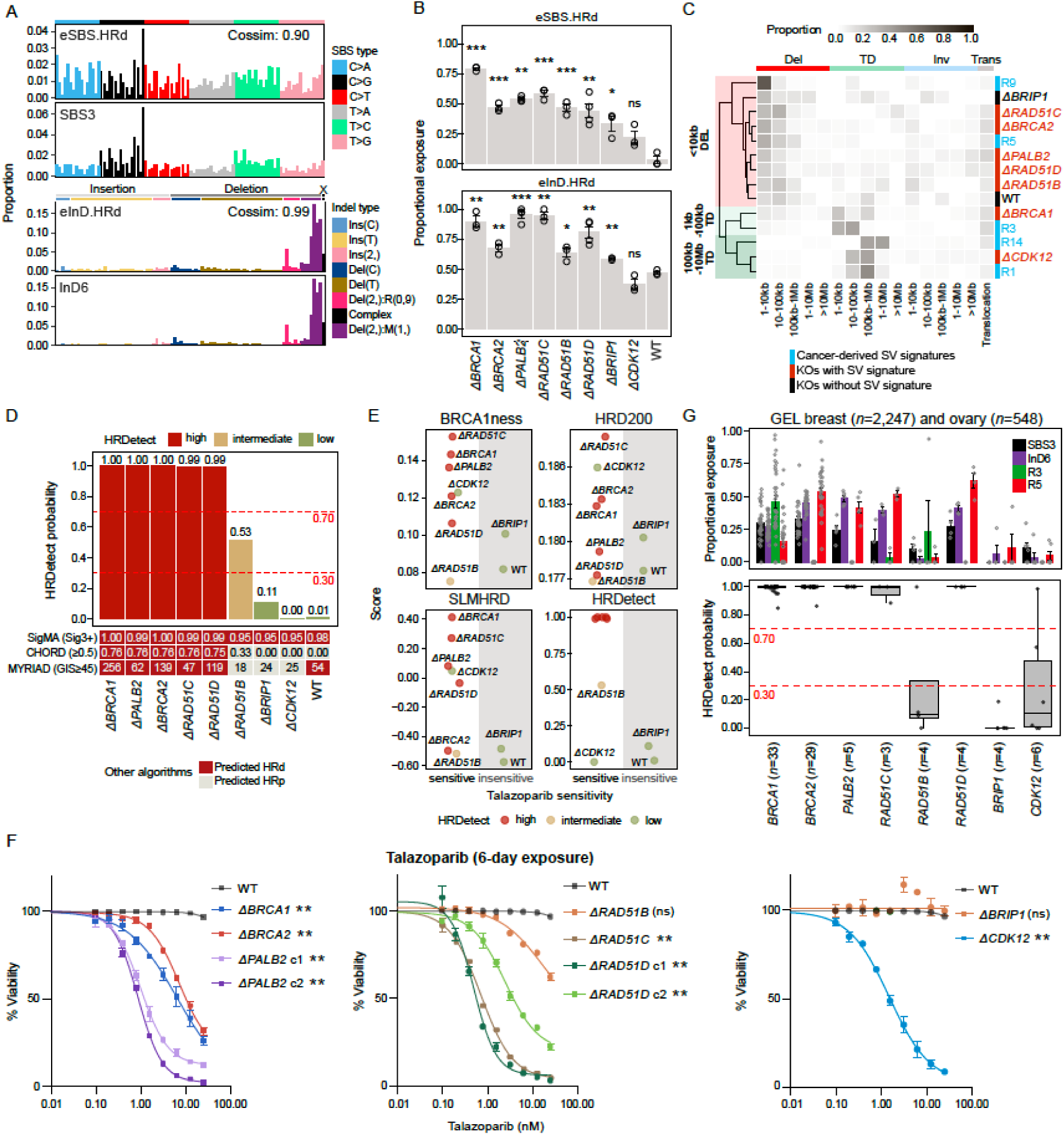
Genomic biomarkers and transcriptomic signatures in predicting HRd and PARPi sensitivities (A) Experimental HRd signatures (eSBS.HRd, eInD.HRd) averaged from genotype-specific signatures in Figure 1D compared to cancer-derived SBS3 and InD6 (RefSig v2, see also **Tables S1, S2, and S3**). Cosine similarities (Cossim) are shown. (B) Proportional exposure of eSBS.HRd and eInD.HRd across RPE1 isogenic knockouts (**Tables S4 and S5**). Error bars represent mean□±□SEM. Two-tailed Welch’s t test versus WT; *p*≤0.05 (*), *p*≤0.01 (**), *p*≤0.001 (***). (C) Unsupervised hierarchical clustering of genotype-specific rearrangement profiles and cancer-derived rearrangement signatures (**Table S6**). (D) HRd prediction by different clinical classifiers across RPE1 isogenic knockouts, ordered by HRDetect score (**Table S7**). Horizontal dashed lines represent thresholds distinguishing HRDetect groups. GIS, genome instability score. (E) Comparison of published transcriptomic HRd biomarker tools (BRCA1ness^31^, HRD200^29^, and SLM^HRD32^) versus HRDetect in predicting PARPi sensitivity (**Table S9,** see also panel G). “Scores” on *y*-axis report degree of HR deficiency based on each classifier. Established thresholds are as follows: BRCA1ness, >-0.3; HRD200, >0.5; HRDetect, >0.7. There was no proposed threshold for SLM^HRD^. (F) Dose response of RPE1 isogenic knockouts to PARPi talazoparib. *n*=3 technical replicates. Error bars represent mean□±□SEM. Two-sided Wilcoxon test (knockout versus WT); *p*≤0.05 (*), *p*≤0.01 (**). (G) Prevalence of HRd signatures and HRDetect prediction in GEL breast (*n*=2,247) and ovarian (*n*=548) cancer cohorts, stratified by HR gene. Top: Proportional exposures of SBS3, InD6, and rearrangement signatures R3 and R5 in samples with biallelic HR gene mutations. Error bars represent mean□±□SEM. Bottom: HRDetect probabilities by HR driver gene. Horizontal dashed lines represent thresholds distinguishing HRDetect groups. See also Figures S2 and S3; Tables S1-S7, S9.

Beyond canonical HR genotypes, RAD51 paralogs (*ΔRAD51B*, *ΔRAD51C*, *ΔRAD51D*), *ΔBRIP1,* and *ΔCDK12* demonstrated increased exposures of eSBS.HRd compared to WT (**Figures 2B and S2A; Tables S2 and S4**). However, *ΔCDK12* did not show increased proportions of eInD.HRd and had an indel profile indistinguishable from WT. *ΔRAD51B* and *ΔBRIP1* exhibited different indel profiles from canonical HRd genotypes (**Figures 2B, S2B and S3C; Tables S3 and S5**). To evaluate rearrangement phenotypes, we performed unsupervised clustering of rearrangement profiles of all genotypes alongside relevant cancer-derived rearrangement signatures^25^. This revealed two main clusters: one dominated by deletions, the other by TDs (**Figures 2C and S2C; Table S6**). *ΔBRCA2*, *ΔPALB2,* and all *ΔRAD51* paralogs clustered with R5 short deletions (<10kb), while *ΔBRCA1* clustered loosely with R3 short TDs (1-10kb). *ΔCDK12* exhibited long tandem duplications (predominantly 100kb-1Mb), in line with R1/14 long TD signatures^25^.

These results provide experimental confirmation that first, typical HRd signatures are not limited to *BRCA1* or *BRCA2* loss and extend to *PALB2*, *RAD51C,* and *RAD51D* defects. Second, mutational heterogeneity exists amongst defective genes in the HR pathway. Third, structural variation (SV) patterns underscore this heterogeneity, with *BRCA1*- and *CDK12*-deficient cells showing distinct tandem duplication biases, while *BRCA2*-like genotypes were enriched for deletions. Together, these findings demonstrate that HRd at the genomic level exists along a continuum – spanning fully penetrant, canonical-like mutational patterns to partial phenotypes, with implications for biomarker development and therapeutic stratification.

### Sensitivity and specificity of HRd classifiers

Given the clinical importance of accurately discerning HRd tumors, we assessed how existing genomic scar-based classifiers performed in our isogenic models. These included the FDA-approved Myriad myChoice HRD score, SigMA^26^ (uses solely SBS3), CHORD^27^ (primarily relies on microhomology-mediated deletions, mh-dels), and HRDetect^4^ (integrates SBS3, SBS8, R3, R5, with genomic features including proportion of mh-dels and HRD index). HRDetect and CHORD correctly identified canonical HR genes and *RAD51* paralog KO genotypes as HRDetect-high (≥0.70), and *ΔBRIP1* and *ΔCDK12* as HRDetect-low (<0.30); both algorithms produced fewer apparent false positives than Myriad and SigMA (**Figure 2D; Table S7**). Notably, *ΔRAD51B* had an intermediate HRDetect score (0.53), consistent with its weaker indel and rearrangement signatures (**Figures 2B, 2D, S2B, S2C, and S3C**). The intermediate HRd status of *ΔRAD51B* was corroborated in two independent *ΔRAD51B* conditional KOs in MCF10A^28^, where eSBS.HRd, eInD.HRd, and R5 signatures were also observed (**Figures S3A and S3B**). The predicted intermediate HRDetect scores aligned with prior literature showing *ΔRAD51B* as partially HR-deficient and as the least sensitive to DNA-damaging agents among KOs of *RAD51* paralogs in U2OS cells^28^. CHORD and Myriad were unable to define intermediate HRd cases. Moreover, SigMA classified all KO genotypes as HR-deficient, emphasizing the limited specificity of using SBS3 as a standalone biomarker for HRd.

Importantly, HRDetect scores generally tracked with PARPi sensitivity and were consistent with HR reporter assay assessment (**Figures 2E and 2F**). HRDetect-high genotypes showed the greatest sensitivity to the clinically used PARPi talazoparib; HRDetect-intermediate Δ*RAD51B* exhibited moderate sensitivity, while HRDetect-low Δ*BRIP1* and WT cells were largely insensitive. Δ*CDK12* was an important exception – despite being HRDetect-low, it was highly sensitive to talazoparib, highlighting that PARPi sensitivity can occur even in the absence of canonical HRd genomic scars.

We further evaluated whether gene expression profiles could serve as reliable biomarkers of HRd. Whole-transcriptome profiles of RPE1 isogenic clones failed to distinguish HRDetect-high from HRDetect-low genotypes (**Table S8**). Restricting the analysis to published transcriptomic HRd signatures^29–31^ or biologically defined synthetic-lethal gene sets for HRd (SLM^HRD^)^32^ did not improve discrimination (**Figure 2E, Table S9**). More importantly, transcriptomic biomarkers showed limited association with PARPi response (**Table S9**). Collectively, these findings underscore the limited discriminatory performance and poor reproducibility of transcriptomic HRd biomarkers. In contrast, composite genomic biomarkers – particularly those integrating multiple mutational features – outperformed transcriptomic signatures in capturing the full spectrum of functional HR impairment (including intermediate states) and showing stronger associations with PARPi sensitivity.

To extend our experimental findings to clinical settings, we analyzed tumors with biallelic *loss-of-function* mutations in HR and related genes within Genomics England 100,000 Genomes Project (GEL 100kGP) breast and ovarian cancer cohorts. Most cases harbored *BRCA1/BRCA2* mutations (*n=*62), followed by *CDK12* (*n*=6) and *PALB2* (*n*=5) (**Figure S3D**). HRd cancers recapitulated the mutational signatures observed in our experimental models, with comparable HRDetect outputs (**Figure 2G**). Tumors with biallelic *RAD51B, CDK12*, or *BRIP1* mutations displayed lower HRDetect scores, consistent with the experimental models. Singular *CDK12^mut^* (GEL-2197374-11) or *RAD51B^mut^* (GEL-2307631-11) cases with HRDetect-high scores were noted, which possibly reflects alternative drivers of HRd-related signatures operating in these two cases.

### HRd genotypes exhibit variable penetrance of synthetic lethal effects

Next, to investigate whether distinct HR gene KOs differ in synthetic lethal (SL) relationships, we used genome-wide CRISPR dropout screens to systematically map the genetic dependencies for each HRd genotype across the isogenic series relative to parental WT cells (**Figure 3A**). This allowed us to identify both negative (*i.e.,* SL effects) and positive genetic interactions (*i.e.,* genetic masking or synthetic rescue (SR) effects) (**Figures 3B, S4, and S5; Table S10**).

**Figure 3.**
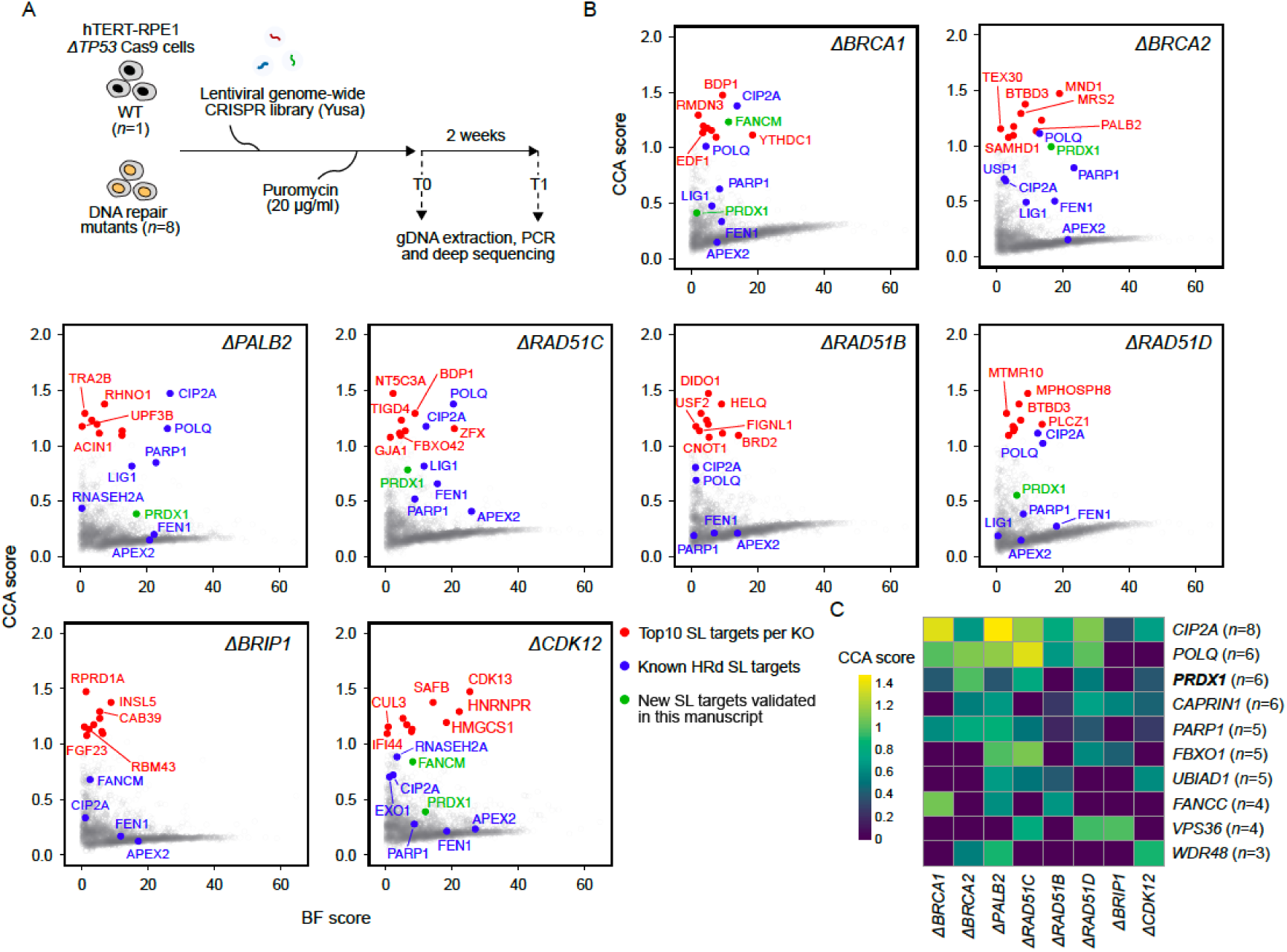
Genome-wide CRISPR screens reveal shared and genotype-specific synthetic lethal interactions (A) Schematic of genome-wide CRISPR dropout screens in isogenic hTERT-RPE1 Δ*TP53* (WT) and eight HR gene knockout genotypes to identify genotype-specific synthetic lethal interactions. (B) Scatter plot of BAGEL2^80^ Bayes factors (BF scores, *x* axis) versus CRISPR Count Analysis^33^ (CCA) scores (*y* axis) for each genotype (**Table S10**). (C) Top ten most common SL targets shared ≥3 HRd genotypes. *n* denotes the number of genotypes in which the target gene is synthetically lethal (CCA≥0.3). See also Figures S4, S5, S6, and S7; Table S10.

Several well-characterized *BRCA1/BRCA2* SL effects showed broad but variable penetrance across the isogenic series (**Figures 3C and S6**). For example, *CIP2A*, which supports mitotic DNA integrity and fitness in *BRCA1-/BRCA2-* mutant cells^33^, was the most consistent and penetrant SL target across all HRd genotypes. *PARP1*^34,35^ and *POLQ*^36,37^ – the drug targets of either approved (PARP1 inhibitors) or in clinical development (POLQ inhibitors) HRd-targeted therapies, showed strong SL effects in *ΔBRCA1, ΔBRCA2, ΔPALB2, ΔRAD51C*, and *ΔRAD51D* cells (aligning with prior functional and clinical observations), but not in *ΔBRIP1*^38^. We confirmed that although *ΔBRIP1* exhibited the expected sensitivity of HRd cells to cisplatin, it showed minimal sensitivity to PARPi, with no detectable response to talazoparib and only modest sensitivity to olaparib (**Figures 2F and S7A**). *PARP1* also failed to reach our SL screen threshold in *ΔCDK12* (**Figures 3B, 3C, and S6**). However, in validation experiments, *ΔCDK12* was selectively sensitive to both talazoparib and olaparib (consistent with prior observations^22,39,40^), despite lacking canonical HRd signatures (**Figures 2F and S7A**).

Other reported *BRCA1/BRCA2* SL targets exhibited even greater genotype-specific variation. Dependencies involving *FEN1, APEX2*^41^, *USP1*^42^, and *LIG1*^43^ varied substantially across HRd genotypes, with strong effects in some contexts but weak or absent effects in others (**Figures 3B and S5**). This variable penetrance of SL effects was independently confirmed using small-molecule inhibitors targeting either USP1 or FEN1 (**Figures S7B and S7C**). Other HRd genotypes, notably *ΔCDK12,* displayed a distinct SL profile consistent with its known biology: the most significant *CDK12* SL partner was its paralog *CDK13* – a transcriptional cyclin-dependent kinase recently validated as a selective vulnerability in *CDK12*-mutant tumors^40^. Additional *CDK12* SL partners were enriched in genes involved in transcriptional elongation and RNA processing, consistent with CDK12’s role in regulating RNA polymerase II^44,45^ (**Figure S7D**).

Beyond known targets, our screens also uncovered new SL dependencies shared amongst HRd genotypes (**Table S10**). These include *NSMCE3* (also known as *NDNL2*) and *STAG2*, which encode proteins involved in sister chromatid cohesion; *CAPRIN1*, which encodes an RNA-binding protein involved in cell cycle regulation and apoptosis^46^; *HMCES*, a gene encoding a sensor of abasic DNA lesions^47^; *FBXO4*, a member of the SCF E3 ubiquitin ligase complex; *WDR48*, which encodes a serine-threonine phosphatase, recently found to be involved in SL interactions with *USP1/LIG1/FEN1*^48^; and *PRDX1* (Peroxiredoxin 1), a gene previously identified as synthetically lethal with *ATM*^49^. Following *CIP2A* and *POLQ, PRDX1* emerged as the third most common SL target across multiple HRd genotypes (**Figure 3C**). When we re-analyzed published genome-wide CRISPR screens that contain a modest number of *BRCA1/2* mutant tumor cell lines^51^, we found *PRDX1* synthetic lethality was also evident here (**Figure 4A**). Orthogonal validation using siRNA and a small-molecule PRDX1 inhibitor confirmed the PRDX1:HRd SL relationship (**Figures 4B, 4C, and S8A**). PRDX1 is an antioxidant enzyme that mitigates intracellular reactive oxygen species (ROS) accumulation in response to DNA damage^50^. Consistent with this function, *PRDX1* loss elevated baseline intracellular ROS levels (**Figure 4D**) and both baseline and DNA damage-induced γH2AX levels (**Figures 4E and S8B**), whereas the antioxidant N-acetylcysteine (NAC) completely reversed PRDX1 inhibitor cytotoxicity in *ΔBRCA1* cells (**Figures 4F and 4G**). In the absence of exogenous DNA damage, the fraction of cells with nuclear RAD51 foci was higher in the absence of *PRDX1* (**Figures 4H and S8C**), suggesting that *PRDX1* loss causes an increase in DNA lesions normally repaired by HR. To understand what these lesions might be, we used a 45-day mutation accumulation assay and found that *ΔPRDX1* subclones acquired an excess of C>A mutations characteristic of oxidative damage-associated SBS18 (**Figures 4I and 4J; Table S2**). It thus seems possible that in the absence of *PRDX1*, the basal burden of oxidative DNA damage increases to an extent that these ultimately form lesions best repaired by RAD51-mediated processes; in the absence of HR, the loss of *PRDX1* thus causes synthetic lethality.

**Figure 4.**
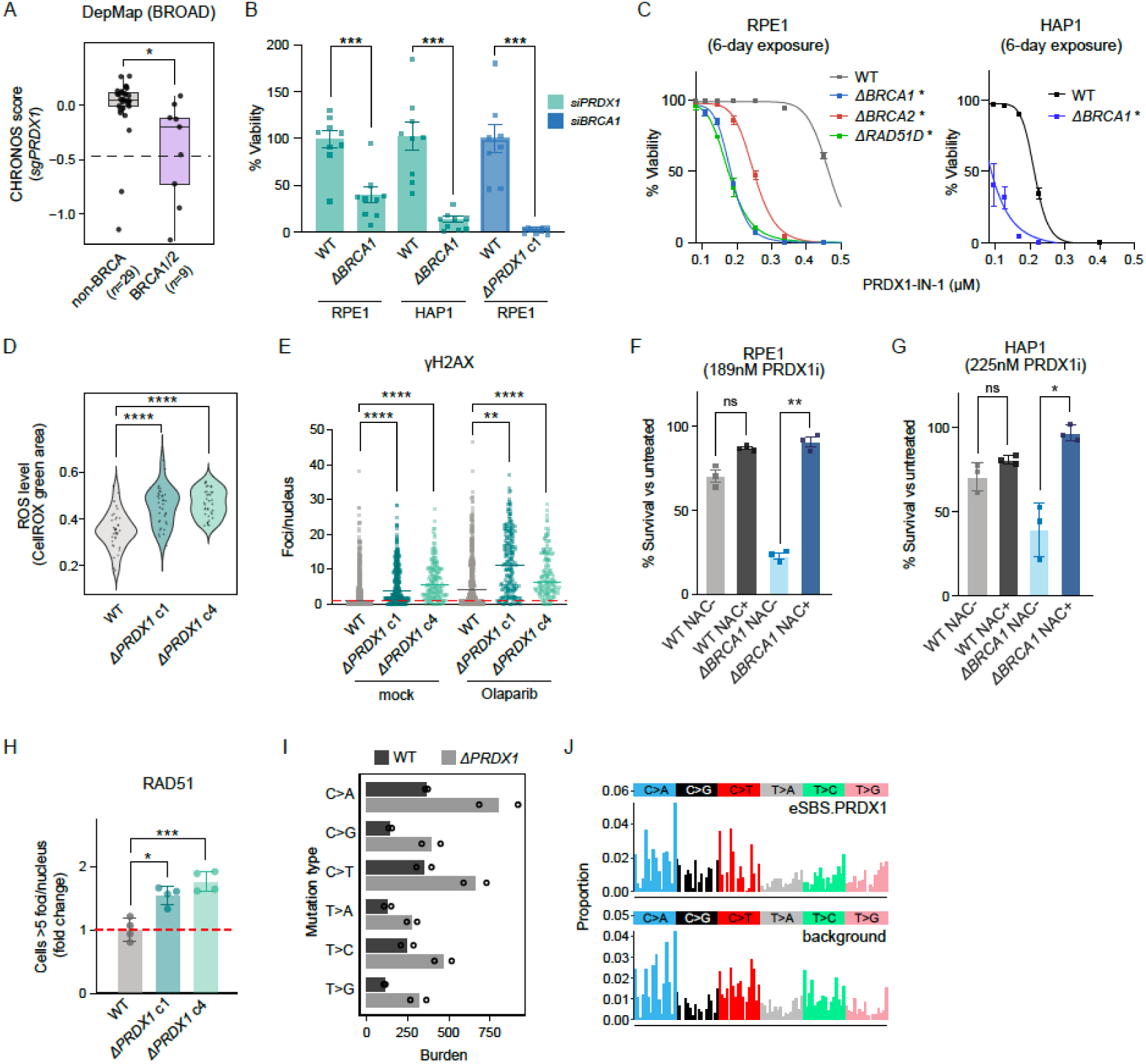
PRDX1 is a ROS-dependent synthetic lethal target in HRd cells (A) *PRDX1* essentiality (CHRONOS scores from DepMap) in biallelically inactivated *BRCA1/2*-mutant (*n*=9) and PARPi-insensitive non-*BRCA* mutant cell lines (*n*=29). Horizontal dashed line marks essentiality threshold. Two-sided Welch’s t-test; *p*≤0.05 (*). (B) Viability following siRNA knockdown of *PRDX1* in RPE1 and HAP1 Δ*BRCA1* cells, and *BRCA1* knockdown in RPE1 Δ*PRDX1* cells. (C) Dose response to PRDX1 inhibitor (PRDX1-IN-1) in HRd isogenic lines (RPE1, HAP1). *n*=3 replicates. Error bars represent mean ± SEM. Two-tailed Wilcoxon test versus respective WT; *p*≤0.05 (*). (D) Baseline reactive oxygen species (ROS) levels quantified with CellROX Green in RPE1 WT and Δ*PRDX1* cells. *n*=36 technical replicates. Two-sided Welch’s t test; *p*≤0.05 (*), *p*≤0.01 (**), *p*≤0.001 (***), *p*≤0.0001 (****). (E) γH2AX foci per nucleus in RPE1 WT and Δ*PRDX1* cells ± 24h olaparib exposure. Center lines represent medians. Horizontal dashed line marks WT median. Two-way ANOVA; *p*≤0.05 (*), *p*≤0.01 (**), *p*≤0.001 (***), *p*≤0.0001 (****). Related to Figure S8B. (F) RPE1 WT and Δ*BRCA1* survival upon 189nM PRDX1i (PRDX1-IN-1) ± N-acetylcysteine (NAC) treatment for 6 days. % viability normalized to mock treated condition per genotype and treatment arm. *n*=3 replicates. Error bars represent mean ± SEM. Two-tailed paired t test; *p*≤0.05 (*), *p*≤0.01 (**). (G) As in (F), for HAP1 cells using 225 nM PRDX1i. (H) Fold change in cells with ≥5 RAD51 foci per nucleus in RPE1 WT and Δ*PRDX1* cells. Error bars represent mean ± SEM. One-way ANOVA; *p*≤0.05 (*), *p*≤0.01 (**), *p*≤0.001 (***). Horizontal line marks the WT mean. Related to Figure S8C. (I) SBS mutation burdens in RPE1 Δ*PRDX1* c1 versus WT. Bar represents mean from *n*=2 subclones per genotype. (J) Background-subtracted SBS signature associated with Δ*PRDX1* (eSBS.PRDX1) and WT background profile. See also Figure S8.

Our screens also indicated that synthetic rescue (SR) effects (*i.e.,* where a gene defect causes less loss of fitness in HRd knockout cells compared to WT cells) also varied according to HRd genotype (**Figure S5**). Consistent with known *BRCA1* epistasis, *ΔBRCA1* cells displayed strong and selective SR effects with *TP53BP1,* the Shieldin complex subunit genes *SHLD1* (also known as *C20orf196*), and *SHLD2* (also known as *FAM35A*), as well as with *CTC1, STN1*, and *TEN1*, components of the CST complex^16,33,51^. These observations were consistent with prior reports showing that loss of these genes partially restores DNA end-resection, rescues HR defects and fitness in *BRCA1*-, but not *BRCA2-*deficient cells^52,53^. Similarly, *ΔRAD51C* and *ΔRAD51D* cells were resistant to the deleterious effects on fitness caused by targeting *RAD51*, *RAD51C,* or *RAD51D*, as well as *RTEL, XRCC2, XRCC3*, *ERCC4,* or *FANCI*. These SR effects were consistent with epistatic relationships within the RAD51 paralog network and reflect pathway-level buffering when components of the same repair module are simultaneously perturbed. Consistent with the downstream role of *BRIP1* (or *FANCJ*) in the Fanconi Anemia (FA)/HR pathway, *ΔBRIP1* cells demonstrated SR interactions with *FANCD2* and *FANCI,* the two components of the FA ID2 complex^54^. Collectively, these expected epistatic relationships provided an internal validation of screen fidelity and supported interpretation of the genotype-specific SL dependencies described above.

### *BRCA1-*specific SL relationships with Fanconi anemia pathway genes

Our isogenic series also uncovered strong SL interactions in *ΔBRCA1* with a broad spectrum of FA pathway genes, including *FANCA, FANCB, FANCC, FANCD2*^55^*, FANCE, FANCF, FANCG, FANCL, FANCI, FANCM,* and *FAAP24* (also known as *C19orf40*) **(Figure S6)**. In contrast, FA genes showed little or no SL effect in *ΔBRCA2*, except for *FANCN* (also known as *PALB2*) alongside its interacting E3 ubiquitin ligase *RNF168* (**Figure 5A**).

**Figure 5.**
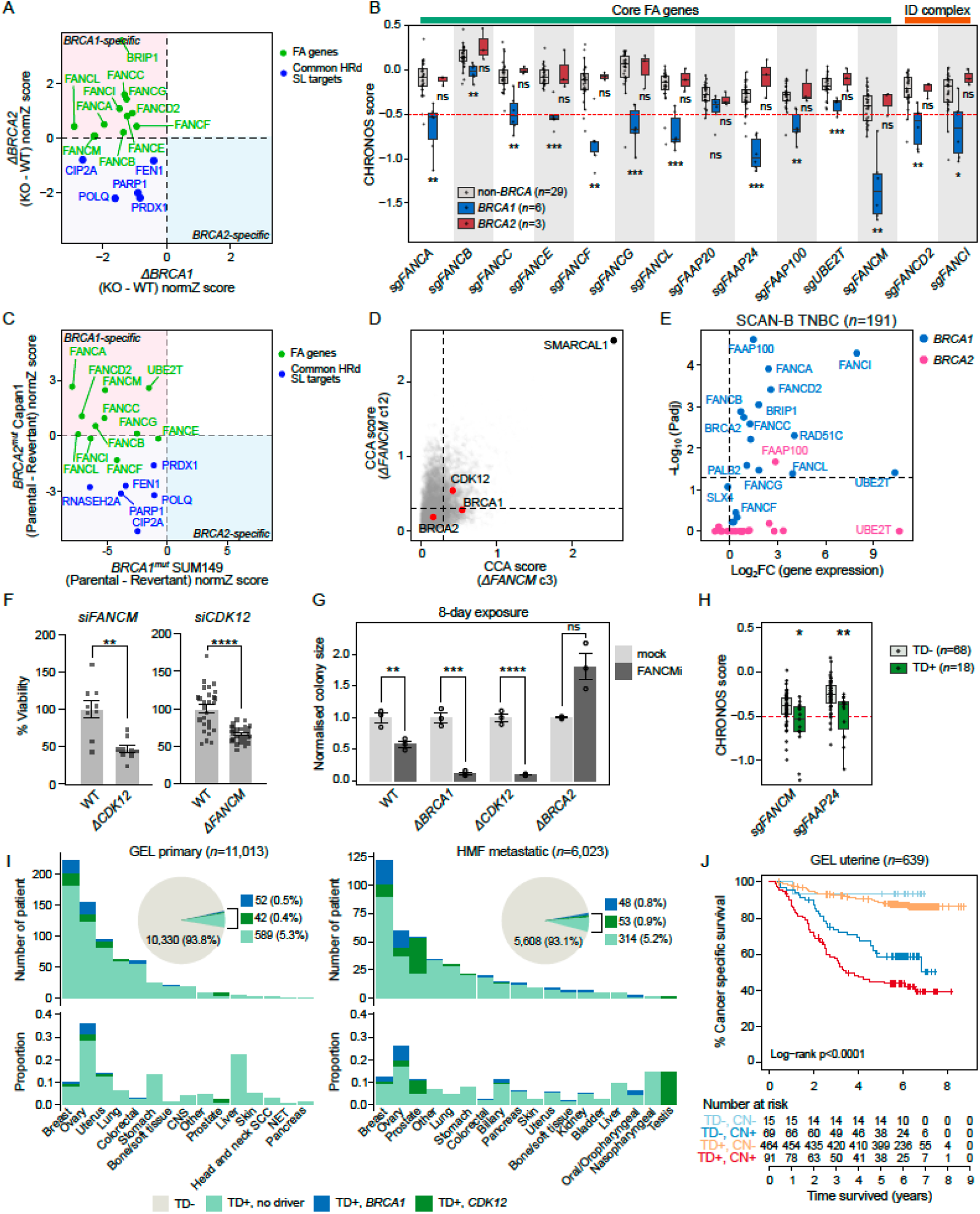
*FANCM* dependency is selective in TD-positive genotypes (*BRCA1, CDK12*) but not *BRCA2*. (A) Scatter plot illustrating differential viability scores (normalized *z*-score, normZ) of Fanconi anemia (FA) pathway genes and known HRd SL targets in isogenic RPE1 Δ*BRCA1*–WT and Δ*BRCA2*–WT CRISPR screens. (B) Boxplots of essentiality scores (CHRONOS) for the indicated genes derived from the BROAD DepMap Project (https://depmap.org/). Cell lines were grouped according to whether they harbored *BRCA1* or *BRCA2* biallelic inactivating mutations (*n*=6, *n*=3, respectively) or not (*n*=29). Similar to Figure 4A. Horizontal red dashed line marks essentiality threshold. (C) As in (A), for *BRCA2^mut^* Capan1 versus *BRCA1^mut^* SUM149 parental-revertant pair isogenic CRISPR screens^32^. (D) Scatterplot of CCA scores for published CRISPR synthetic lethality screen^57^ in RPE1 Δ*FANCM* (clones c3 and c12), highlighting *BRCA1, CDK12*, and *BRCA2* against *SMARCAL1* (a known profound SL target to *FANCM*). (E) Differential expression of FA genes in SCAN-B TNBC samples (*n*=191), contrasting *BRCA1-* and *BRCA2-* cancers respectively, to samples without HR gene mutations and are HRDetect-low. (F) Viability of RPE1 cells following reciprocal siRNA knockdown: *FANCM* in Δ*CDK12* and *CDK12* in Δ*FANCM*. *n*=3 replicates. Error bars represent mean ± SEM, two-tailed Welch’s t test versus respective WT; *p*≤0.05 (*), *p*≤0.01 (**), *p*≤0.001 (***), *p*≤0.0001 (****). (G) Fold change in colony size of RPE1 WT, Δ*BRCA1,* Δ*CDK12,* and Δ*BRCA2* cells treated with FANCM inhibitor (FANCM-BTR PPI-IN-1, 25μM for 8 days) versus mock treatment. Each genotype was normalized to its corresponding mock condition. *n*=3 replicates. Error bars represent mean ± SEM, two-tailed Welch’s t test compared to the respective mock condition per genotype; *p*≤0.05 (*), *p*≤0.01 (**). Related to Figures S8E and S8F. (H) CHRONOS essentiality scores of *FANCM* and *FAAP24* derived from the BROAD DepMap project. Cell lines were fitted with rearrangement signatures and grouped according to whether they harbored TD signatures (R1, R3, R14) (TD+, *n*=18) or not (TD-, *n*=68). Two-sided Welch’s t-tests; *p*≤0.05 (*), *p*≤0.01 (**). (I) Prevalence of samples with TD signatures with or without *CDK12* and *BRCA1* biallelic *loss-of-function* mutations in primary (GEL, *n*=11,013) and metastatic (HMF, *n*=6,023) pan-cancer cohorts (**Table S11**). (J) Kaplan–Meier analysis of cancer-specific survival in GEL uterine cancer cohort (*n*=639), stratified by tandem duplication status (TD+/TD−) and copy-number status (CN+/CN−). Samples of unknown copy number status were removed. The *p* value was calculated using two-sided log-rank test to compare survival between groups. See also Figures S8 and S9; Tables S11.

To validate the *BRCA1*-specific general FA dependency, we interrogated three independent datasets. First, analysis of the DepMap^56^ CRISPR dependency data confirmed that FA gene dependencies were selectively enriched in *BRCA1*- and not *BRCA2-* mutant lines (**Figure 5B**). Among FA genes, *FANCM* displayed the most pronounced genotype-selective SL effect. Second, reanalysis of published genome-wide CRISPR screens^32^ in *BRCA1^mut^* SUM149 and *BRCA2*^mut^ Capan1 parental-revertant pairs showed that core FA genes exhibited *BRCA1*-specific SL, whereas canonical HRd SL targets such as *PARP1*, *POLQ*, and *CIP2A* were shared between both genotypes (**Figure 5C**). Third, re-examination of published hTERT-RPE1 Δ*FANCM* screens^57^ also revealed a stronger SL effect with *BRCA1* than with *BRCA2*, consistent with our results (**Figure 5D**).

Transcriptomic buffering, where the loss of a tumor suppressor gene is accompanied by compensatory overexpression of its SL partners, is a common phenomenon across cancers^32^. Using this framework, we examined expression profiles of FA pathway genes in triple-negative breast cancers (TNBCs), stratified by HR gene mutations^58^. Among HRd cases, *FANCI*, *FAAP100*, *FANCA*, and *FANCD2* were significantly upregulated in biallelically mutated *BRCA1,* but not *BRCA2* cancers (**Figure 5E**), consistent with the idea that upregulated FA genes in *BRCA1*-deficient tumors preserve cellular fitness, thereby creating a *ΔBRCA1* genotype-selective vulnerability to FA gene disruption.

### TD phenotype as a biomarker for FANCM dependency

In our screen, *FANCM* also emerged as a prominent SL target in Δ*CDK12*, in addition to Δ*BRCA1* (**Figure 3B**). *FANCM* has been proposed as a therapeutic target based on its SL interactions with *BRCA1*, *SMARCAL1*, *RAD52*^48,57,59^ and its essentiality in tumor cells that utilize the alternative lengthening of telomeres (ALT) pathway^60,61^. While current drug development efforts focus on ALT-positive tumors, our systematic study reveals that *FANCM*-dependent genotypes (*i.e.,* Δ*BRCA1* and Δ*CDK12*) have TD signatures in common. This raises the possibility that TD phenotype, in addition to specific driver mutations, might serve as a biomarker for FANCM-targetable tumors beyond ALT-dependent cancers^61^.

We first validated *FANCM:CDK12* SL using siRNA, demonstrating the reciprocal loss of viability following depletion of *FANCM* in Δ*CDK12* cells and *CDK12* in Δ*FANCM* cells (**Figures 5F and S8D**). Pharmacological disruption of FANCM using a FANCMi (FANCM-BTR-IN-1)^62^ phenocopied this SL interaction, selectively impairing the viability of Δ*BRCA1* and Δ*CDK12*, while having a more modest effect in WT and Δ*BRCA2* cells (**Figures 5G, S8E, and S8F)**. Analysis of WGS data from DepMap cell lines^56,63^ revealed that cancer cell lines harboring TD signatures (R1, R3, R14) exhibited significantly stronger dependencies on *FANCM* and its interacting partner *FAAP24* (lower CHRONOS scores) compared to those without TD (**Figure 5H**). This association persisted even after excluding *BRCA1*-deficient (*i.e.,* HRDetect_≥_0.7) and biallelically *CDK12*-mutated lines, though with reduced statistical significance likely due to smaller sample size (**Figures S8G and S8H**). Notably, no analogous relationship was observed for *CDK13*, the paralog SL partner of *CDK12* – supporting the notion that *FANCM* dependency tracks with TD phenotype, rather than being exclusive to *BRCA1*^59^ and/or *CDK12* mutations *per se* (**Figure S8G**).

To evaluate the potential of TD phenotype as a candidate biomarker of FANCM inhibition or general TD-associated dependencies, we examined TD signatures R1, R3, and R14 across 11,013 primary cancers (GEL) and 6,023 metastatic cancers (Hartwig Medical Foundation (HMF)) (**Figure S9A, Table S11**). Notably, the vast majority of TD+ tumors – 86.2% primary (*n*=589) and 75.7% of metastatic (*n*=314) cases – lacked identifiable *BRCA1* or *CDK12* driver mutations (**Figure S9A**). While TD signatures were most prevalent in breast and ovarian cancers, uterine cancer ranked third by the number of TD+ cases among primary cancers, which was notable because copy-number-high (CN+) uterine cancers have been associated with poor clinical outcome (**Figure 5I)**. Consistent with this, TD positivity significantly overlapped with CN+ status in GEL uterine cohort (**Figure S9B**). TD+/CN+ was associated with the poorest overall survival, although this did not reach statistical significance, likely due to limited events after adjustment for multiple clinical covariates (HR=7.42, 95% CI=1.00-55.13, *p*=0.05) (**Figure 5J**). This subgroup of TD+/CN+ patients potentially represents a candidate demographic for evaluating FANCM-targeted strategies. Overall, >5% of all analyzed tumors exhibited TD phenotypes without canonical/identifiable drivers – representing a substantial patient population that could potentially benefit from FANCM-targeted therapy guided by genomic markers rather than genotype alone.

We next sought to characterize the shared cellular and molecular features underlying the repair defect in TD-enriched genotypes. At the cellular level, *FANCM* depletion by CRISPR-Cas9 led to the accumulation of persistent 53BP1 nuclear bodies in Δ*BRCA1* or Δ*CDK12* cells, indicative of unresolved DNA damage carried over from the previous cell cycle, but did not elicit the same phenotype in Δ*BRCA2* or WT cells (**Figures 6A and S9C**). Correspondingly, *FANCM* CRISPR-Cas9 targeting in Δ*BRCA1* or Δ*CDK12* cells caused severe genome instability phenotypes marked by increased formation of micronuclei and γH2AX, whereas Δ*BRCA2* and WT cells were relatively unaffected (**Figures 6B, 6C, S9D, and S9E**). The selective emergence of these chromosomal abnormalities in *BRCA1-* and *CDK12-* deficient backgrounds suggests that FANCM becomes essential for maintaining genome integrity in specific HR-deficient contexts, rather than universally across the full spectrum of HRd.

**Figure 6.**
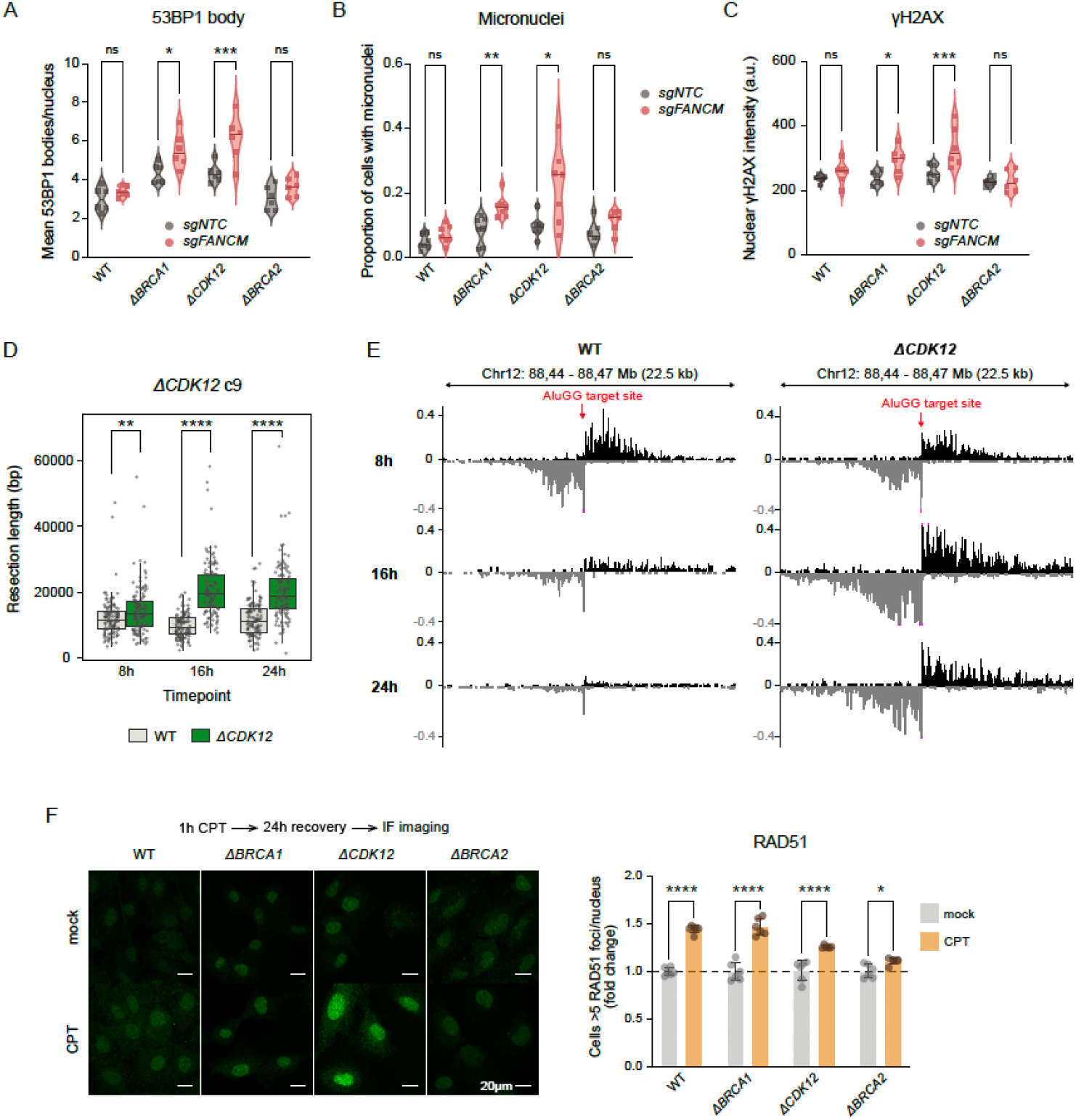
*BRCA1-* and *CDK12-* deficient cells share repair defects distinct from *BRCA2* (A) 53BP1 nuclear bodies per nucleus following *FANCM* depletion. Center lines show the medians. Two-way ANOVA; *p*≤0.05 (*), *p*≤0.01 (**), *p*≤0.001 (***). (B) Proportion of cells with at least one micronucleus. Center lines show the medians. Two-way ANOVA; *p*≤0.05 (*), *p*≤0.01 (**). (C) γH2AX nuclear intensity following *FANCM* depletion. Center lines show the medians. Two-way ANOVA; *p*≤0.05 (*), *p*≤0.01 (**), *p*≤0.001 (***). (D) END-seq resection tract lengths in RPE1 Δ*CDK12* (clone c9) at 8h, 16h, and 24h post G1 release. Welch’s t-tests; *p*≤0.05 (*), *p*≤0.01 (**), *p*≤0.001 (***), *p*≤0.0001 (****). (E) Genome browser screenshots displaying END-seq signals at a representative nick-induced DSB generated by nCas9(D10A) and sgAluGG in RPE1 WT and Δ*CDK12.* Arrow indicates AluGG target site where nick occurs. The accumulated reads flanking the target site indicate end resection. Reads in black and gray represent plus and minus strands, respectively. Cells were collected 8h, 16h, and 24h post G1 release. (F) Quantification (left) and representative immunofluorescence images (right) of RAD51 foci in RPE1 WT, Δ*BRCA1*, Δ*CDK12,* and Δ*BRCA2* post 1h of 1μM CPT exposure and 24h recovery. Error bars represent mean ± SEM. One-way ANOVA; *p*≤0.05 (*), *p*≤0.01 (**), *p*≤0.001 (***), *p*≤0.0001 (****). See also Figure S9.

Given FANCM’s established role in replication fork remodeling and repair pathway choice at stalled forks^59,64,65^, we next examined replication-associated DSB repair in these genotypes. Using END-seq to monitor repair of nick-induced DNA breaks^66^, we found that end resection was intact in Δ*CDK12* cells; median resection tract lengths were, in fact, modestly increased in Δ*CDK12* compared to WT, consistent with prior observations in *BRCA1-null* cells^66^ (**Figures 6D and 6E**). Nevertheless, despite competent resection, Δ*CDK12* cells exhibited delayed and defective repair of nick-induced double-strand breaks, indicating a repair defect downstream of resection (**Figure 6E**), like Δ*BRCA1*. We hypothesize that Δ*BRCA1* and Δ*CDK12* cells retain limited capacity to load RAD51 at replication-coupled DSBs, resulting in attenuated RAD51 engagement^66^. In contrast, *BRCA2* loss abrogates RAD51 nucleation altogether, because BRCA2 is required to overcome the inhibitory barrier of RPA and directly load and stabilize RAD51 on RPA-coated ssDNA^67^. We therefore examined RAD51 engagement following camptothecin (CPT) treatment, which induces replication-dependent double-strand breaks^68^, and found that Δ*BRCA1* and Δ*CDK12* cells were able to form RAD51 foci, whereas Δ*BRCA2* cells failed to form RAD51 foci altogether (**Figures 6F, S9F, and S9G**).

Although the precise molecular events linking attenuated RAD51 engagement to TD formation, and how FANCM functions in this process, remain to be fully defined, our data indicate that *BRCA1-* and *CDK12-*deficient cells converge on a shared replication-associated repair defect (downstream of resection) characterized by attenuated RAD51 engagement – distinct from the RAD51-absent state in *BRCA2* deficiency. Prior work has shown that FANCM limits TD formation, and that *BRCA1* loss is synthetic lethal with FANCM ATPase inactivation, with evidence for synergistic TD induction when *BRCA1* and *FANCM* are both compromised^59^. We therefore propose that FANCM dependency in TD-positive cells reflects a heightened reliance on FANCM-mediated stalled fork processing and repair when RAD51 engagement is attenuated but not absent. In this context, FANCM becomes essential to prevent the accumulation of toxic repair intermediates and consequent genome instability.

### HRd genotypes influence PARPi resistance mechanisms

Finally, we asked whether distinct HRd genotypes differ in their routes to PARPi resistance. Beyond genetic reversion of the mutated HR genes (as has been recurrently seen for *BRCA1, BRCA2, PALB2, RAD51C and RAD51D*), most of what is known about PARPi resistance has emerged from preclinical studies in *BRCA1*-mutant cells^17,69^. These studies have shown that loss of *TP53BP1, DYNLL1,* or the Shieldin complex causes PARPi resistance in *BRCA1-* but not *BRCA2-* mutant cells or cancers^18,70^. Far less is known about how PARPi resistance might emerge in *BRCA2*-mutant cancers^69^. To address this, we performed PARPi CRISPR–Cas9 resistance screens in the isogenic Δ*BRCA1* and Δ*BRCA2* RPE1 cells (**Figures 7A and S10A; Table S12**), which revealed both shared and genotype-restricted resistance mechanisms, indicating that the molecular routes to PARPi resistance are not uniform across HR-deficient states. For example, *E2F7* loss – a known mechanism of PARPi resistance caused by de-repression of RAD51 transcription^71,72^ – was observed in both Δ*BRCA1* and Δ*BRCA2* cells; whereas, as expected, loss of *TP53BP1*, *DYNLL1*, or *SHLD1* caused PARPi resistance specifically in Δ*BRCA1* cells (**Figure 7A**). Conversely, we identified several resistance drivers more prominent in Δ*BRCA2* than in Δ*BRCA1* cells, including loss of *BRIP1*^38^ or the helicase *RECQL5.* These genotype-specific findings were replicated in independent screens in *BRCA1^mut^* SUM149 and *BRCA2^mut^*Capan1 cells (**Figures 7B and S10B**). Further assessment confirmed that in RPE1 Δ*BRCA2* cells, CRISPR-Cas9 targeting of *RECQL5* or *BRIP1* significantly increased olaparib resistance, whereas neither had a significant effect in isogenic Δ*BRCA1* cells (**Figures 7C, S10C, and S10D**). Strikingly, CRISPR-Cas9 targeting of BRIP1 further reduced survival of Δ*BRCA1* cells exposed to olaparib. Supporting this, *BRIP1* loss has been shown to reduce PARP1 trapping but also to be synthetically lethal with *BRCA1* dysfunction^38^, providing one explanation as to why its loss causes PARPi resistance in Δ*BRCA2* but lethality in Δ*BRCA1* cells. *RECQL5*-defective cells have been shown to accumulate *BRCA1*-dependent RAD51 foci^73^, perhaps explaining why *RECQL5* loss confers PARPi resistance in *BRCA2-* but not *BRCA1-* deficient cells.

**Figure 7.**
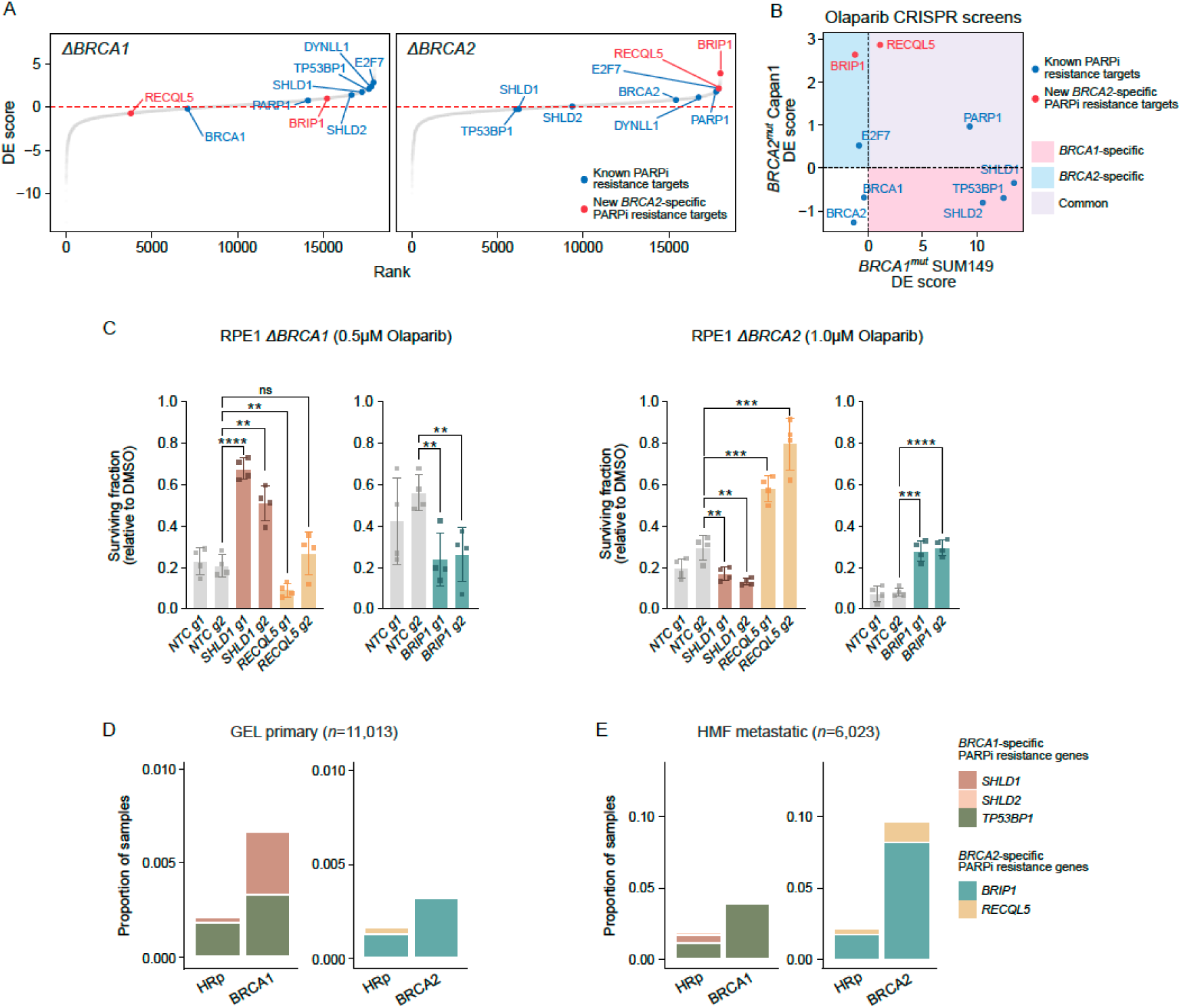
PARPi resistance mechanisms are genotype-specific (A) Drug effect (DE) scores of gene targets from olaparib CRISPR resistance screens in RPE1 Δ*BRCA1* and Δ*BRCA2* cells (**Table S12**). (B) DE scores of known PARPi resistance genes and new *BRCA2*-specific PARPi resistance genes from olaparib CRISPR resistance screens in *BRCA1^mut^* SUM149 and *BRCA2^mut^*Capan1 cells. (C) Viability of RPE1 Δ*BRCA1* (left) and Δ*BRCA2* (right) treated with olaparib, with or without sgRNA-mediated knockout of *SHLD1*, *BRIP1,* or *RECQL5*. Unpaired t test; *p*≤0.05 (*), *p*≤0.01 (**), *p*≤0.001 (***), *p*≤0.0001 (****). (D) Frequency of biallelic *loss-of-function* mutations in *BRCA1*-specific (*i.e., SHLD1, SHLD2, TP53BP1*) and *BRCA2*-specific (*i.e., BRIP1, RECQL5*) resistance genes in primary cancers (GEL, *n*=11,013), stratified by HR-proficient (HRp), *BRCA1*-like, and *BRCA2*-like status. *BRCA1*-like: HRDetect-high with SV3 > SV5; *BRCA2*-like: HRDetect-high with SV5 > SV3. (E) As in (D), for metastatic cancers (HMF, *n*=6,023). See also Figure S10; Table S12.

To assess if these resistance drivers occur clinically, we sought biallelic *loss-of-function* (LoF) mutations in these genotype-specific PARPi resistance genes across GEL primary and Hartwig metastatic cohorts. In both cohorts, *BRCA1/BRCA2*-like cancers harbored more biallelic *LoF* mutations in genotype-specific PARPi resistance genes compared to HR-proficient tumors (**Figures 7D and 7E**). Metastatic tumors showed higher frequencies overall, with ∼10% of *BRCA2*-like tumors in Hartwig harboring resistance gene *LoF* mutations compared to ∼0.3% in GEL. Albeit rarer, *BRCA1*-specific resistance mutations affected ∼4% of *BRCA1*-like tumors in Hartwig and 0.6% in GEL (**Figures 7D and 7E)**. Because *BRIP1* is included in clinical ctDNA sequencing panel^74,75^, we assessed whether deleterious *BRIP1* mutations have been observed in women with germline or somatic *BRCA1-* or *BRCA2-* mutated advanced breast cancer who received olaparib as part of their *standard-of-care* treatment^18^. In this cohort of 47 patients, one *BRCA2^mut^* patient with acquired olaparib resistance had a *BRIP1* mutation detected by ctDNA, suggesting that this might be a clinically relevant resistance mechanism.

Taken together, these data suggest that a subset of *BRCA1-/BRCA2*-mutated cancers could harbor pre-existing resistance driver mutations that may blunt PARPi benefit. Prospective evaluation of these genes as exclusion biomarkers when considering PARPi therapy could help refine patient stratification, avoid ineffective PARPi treatment, and guide alternative treatment strategies.

## DISCUSSION

An isogenic series targeting multiple genes within the same pathway provides a powerful means to compare and contrast the impact of individual genetic changes on a common background. This parallel approach allows for a holistic evaluation of all available data at once, revealing comparative insights that would not be apparent in sequential, pairwise studies.

First, we confirmed the nature of typical mutational signatures associated with *BRCA1*/*2* deficiency (SBS3, InD6, R3, R5) and their presence in *ΔPALB2*, *ΔRAD51C,* and *ΔRAD51D*. Although strongly suspected from large-scale cancer genome analysis, this has never been systematically demonstrated in a matched isogenic human cellular model and thus not all these genetic defects are widely represented in clinical assays to date^76^. Moreover, many of the earliest clinical trials were focused on germline mutation carriers. Yet these isogenic models show that somatically acquired mutations are equally important. *ΔBRIP1* stood out as the exception: despite cisplatin sensitivity, it failed to recapitulate the canonical HRd mutational or functional phenotypes and was PARPi-insensitive. Clinically, *BRIP1* is included in HRd panels, however clinical responses in *BRIP1*-mutant cancers to PARPi have been variable. Our data provide a plausible experimental explanation for this heterogeneity.

Second, comparative analysis across HRd scar- or signature-based methods for tumor classification reveals the drawbacks of currently used clinical assays. Myriad’s false positive rate and the lack of intermediate HRd phenotype reporting by all other assays are a limitation, underscored by the demonstration that signature-based scores track PARPi sensitivity and outperform expression profiles as a biomarker of HRd. We do, however, caution against the premise that any single assay should be the panacea for identifying sensitivity to a single compound. Notably, *CDK12*-mutant cells demonstrate vulnerability to PARPi but do not carry the same signatures as canonical HRd tumors. This is an important point of awareness. However, in a clinical setting where WGS is routinely performed and captures all the heterogeneity reported above (rather than requesting many independent tests), we would be better equipped to identify the full precision intervention potential for each person’s tumour.

Third, we show how genetic SL interactions are not binary categorizations. The penetrance of SL interactions varied considerably across genes. *CIP2A* was universally SL with defects in all eight HR genes, but others (*e.g., PARP1, POLQ, APEX2*) were not. This has implications for how we think about why patients do not respond similarly and provides guidance for identifying patients who may warrant combination therapies beyond a single agent.

Fourth, we identified new SL interactions. *PRDX1* emerged as a recurrent redox-regulated vulnerability across multiple HRd genotypes. Our data support a model in which PRDX1’s function provides a critical buffer against endogenous oxidative damage; when exceeded, cells succumb to accumulated lesions requiring HR-mediated repair. The complete rescue of PRDX1 inhibitor-induced cytotoxicity by NAC confirms ROS as the upstream driver of this cumulative stress. Several PRDX1 inhibitors are in preclinical development^77,78^. *PRDX1* targeting may represent a new therapeutic strategy for HRd cancers.

Fifth, we also uncovered a selective SL relationship between *BRCA1* loss and Fanconi anemia pathway genes and focused our attention on the selective *FANCM* dependency in Δ*BRCA1* and Δ*CDK12 –* genotypes both characterized by TD signatures, albeit of different sizes. We confirmed the *FANCM:CDK12* SL relationship and proposed an alternative candidate patient population that could be evaluated for FANCM-targeted strategies, that would be missed by current targeted approaches. We reasoned that the *FANCM* dependency in Δ*BRCA1* and Δ*CDK12* reflects a TD-permissive repair state for which FANCM has essential control. Our data suggest the shared repair defect in Δ*BRCA1* and Δ*CDK12* is downstream of end resection and differs from Δ*BRCA2* in engagement with RAD51. While the precise molecular steps linking reduced RAD51 engagement, TD generation, and FANCM remain an important open question, prior work has shown that FANCM plays a role in limiting TD formation^59^. A plausible model could be that in Δ*BRCA1* and Δ*CDK12* cells, suboptimal RAD51 loading favors aberrant fork restart/bypass via template-switch mechanisms^79^. FANCM prevents runaway replication restart, suppressing excessive template switching and limiting TD formation^59^. FANCM loss therefore exacerbates template-switch cycles, leading to catastrophic cell death. *BRCA2* loss, in contrast, abolishes RAD51 filament stability, negating this dependency altogether.

Finally, CRISPR resistance screens showed that HRd genotypes employ distinct therapy resistance mechanisms. Our data show that BRIP1 loss can have opposing therapeutic implications depending on HRd genotype: it is synthetically lethal in Δ*BRCA1* cells but promotes PARPi resistance in Δ*BRCA2* cells. Thus, resistance biomarkers should be interpreted in relation to the underlying HRd genotype rather than treated as universal predictors of PARPi response. In the future, genotype-informed resistance profiling could extend beyond Δ*BRCA1* and Δ*BRCA2* to identify patients unlikely to benefit from continued PARPi (or any other drugs), enabling transition to alternative therapies (*e.g.,* POLQ inhibitors).

Our overarching message is that HRd is not simply a single molecular state, and thus not a single clinical entity that can be managed with a uniform strategy. The heterogeneity we described – spanning HRd driver genotypes, signatures, SL dependencies, and resistance mechanisms – demands a more granular dissection for more effective precision medicine activities. This work highlights the importance of using the totality of mutational information to inform clinical decision-making per patient. The spectrum of genetic defects together with signature-based strategies should be used to synthesize a holistic interpretation of each patient’s cancer. This would expand the range of patients that could receive targeted therapies of established therapeutics such as PARPis and for investigational compounds such as FANCMis, repurposed for TD phenotypes that define ∼5% of all primary and metastatic cancers.

## Limitations of the study

We acknowledge several limitations of this study. First, the isogenic system was focused on one cellular model, RPE1. Replication of the full experimental series in another isogenic cellular model was neither cost-effective nor feasible within the scope of this study. However, key findings were validated in additional normal and cancer cellular models where appropriate and cross-checked against publicly available datasets where possible. RPE1 is a non-malignant, hTERT-immortalized epithelial model that has been used extensively by the DNA repair community and provides important insights into mutagenesis and functional effects in pre-malignant cellular contexts. Nevertheless, given that RPE1 is derived from normal, primary retinal pigment epithelial cells, these data underscore how mutational signatures and functional effects of HRd can manifest regardless of cell-type or cellular model. Second, our experimental system modeled complete gene loss, which may not reflect partial disruptions or hypomorphic mutations with subtler phenotypes that may arise in human cancers. Third, the 45-day mutation accumulation window may have limited power for detecting rearrangements, which were modest across all genotypes. Finally, the proposed relationship between attenuated RAD51 recruitment, TD formation, and *FANCM* dependency remains correlative and awaits direct mechanistic investigation. Despite these limitations, our systematic comparative approach has produced a rich resource with important conceptual offerings, shifting our understanding of synthetic lethality from a binary to a continuum model, highlighting potential sources of heterogeneity in clinical responses and *a priori* resistance, calling for modernizing the molecular taxonomy of HRd for precision cancer medicine.

## Resource availability

Data from the National Genomic Research Library (NGRL) used in this research are available within the secure Genomics England Research Environment. Access to NGRL data is restricted to adhere to consent requirements and protect participant privacy. Data used in this research include: WGS and substitution, indel, and rearrangement mutational profiles of *n*=11,013 samples. Access to NGRL data is provided to approved researchers who are members of the Genomics England Research Network, subject to institutional access agreements and research project approval under participant-led governance. For more information on data access, visit: https://www.genomicsengland.co.uk/research (https://doi.org/10.6084/m9.figshare.4530893).

## Acknowledgements

We gratefully acknowledge the participants of the National Genomic Research Library (NGRL), whose contributions made this research possible. Secure access to the NGRL under project ID RR239 was provided by Genomics England, which delivers the NGRL in partnership with NHS England, and is wholly owned by the UK Department of Health and Social Care. The NGRL contains participants’ health data collected by the NHS as part of their care, along with samples and data from their participation in research, for which fully informed consent has been obtained. This includes genomic and clinical data provided through the NHS Genomic Medicine Service, as well as data obtained through research studies, including the 100,000 Genomes Project and the Generation Study, both of which are delivered in partnership with the NHS, and from other research cohorts involving external collaborators.

This publication and the underlying study have been made possible partly based on data that Hartwig Medical Foundation has made available to the study through the Hartwig Medical Database.

## Author contributions

G.C.C.K., S.N-Z., C.J.L. conceived this work. G.C.C.K. and S.J.Z. generated the isogenic knockouts, performed the mutation accumulation, validation, and sequencing experiments with help from S.B. and P.J.T.. S.J.Z. and D.B. analyzed the sequencing data with guidance and input from G.C.C.K., A.D., G.R., Y.M., H.R.D.. R.B. and F.S. performed the CRISPR screens/PARPi resistance screens and validated the targets with S.C.. R.B. and S.J.Z. analyzed the screen data. D.B.K. did all the immunofluorescence and foci experiments. V.T. performed END-seq. G.C.C.K., S.J.Z., S.N-Z., C.J.L. wrote the paper, with input from all the authors. S.N-Z. and C.J.L. supervised this work.

## Declaration of interests (should be identical to the declaration of interest form)

Patents: G.C.C.K., G.R., A.D., H.R.D., and S.N-Z. are inventors on patent applications encompassing the code and intellectual principle of the HRDetect algorithm (PCT/EP2017/060294) (H.R.D., S.N-Z.), clinical use of signatures (PCT/EP2017/060289) (H.R.D., S.N-Z.), clinical predictors (PCT/EP2017/060298) (H.R.D., S.N-Z.), rearrangement signature methods (PCT/EP2017/060279) (S.N-Z.), mutational signatures method (PCT/EP2023/056078) (A.D., S.N-Z), indel signature methods (PCT/EP2024/077959) (G.C.C.K., S.N-Z.) and PRRDetect algorithm (PCT/EP2024/078030) (G.C.C.K., G.R., S.N-Z.). A UK patent application relating to the findings described in this manuscript has been filed: GB2611291.2, “Treatment of Cancer,” filed on 14 May 2026. C.J.L. receives and/or has received research funding from AstraZeneca, Merck KGaA, and Artios; received consultancy, SAB membership, or honoraria payments from Syncona, Sun Pharma, Gerson Lehrman Group, Merck KGaA, Vertex, AstraZeneca, Tango, 3rd Rock, Ono Pharma, Artios, Abingworth, Tesselate, Dark Blue Therapeutics, Pontifax, Astex, Neophore, GlaxoSmithKline, Dawn Bioventures, Blacksmith Medicines, and FoRx; has stock in Tango, Ovibio, Hysplex, Tesselate, and Ariceum. He is also a named inventor on patents describing the use of DNA-repair inhibitors, including ATR inhibitors, and stands to gain from their development and use as part of the ICR ‘Rewards to Inventors’ scheme and also reports benefits from this scheme associated with patents for PARP inhibitors, paid into C.J.L.’s personal account and research accounts at the Institute of Cancer Research. Research in Madalena Tarsounas’s laboratory is supported by Cancer Research UK Program Award (DRCPGM\100001).

## Supplemental information titles and legends Supplemental Information

Document S1. Figures S1–S10. Tables S1-13.

## Tables

### STAR Methods

#### Data and code availability

WGS data have been deposited at <u>EGAD50000002406</u> (DOI:); RNAseq data have been deposited at <u>EGAD50000002336</u> (DOI:); CRISPR screen and drug screen data have been deposited at (DOI:). All original code has been deposited at [repository] and is publicly available at [DOI] as of the date of publication.

#### Additional information

Any additional information required to reanalyze the data reported in this paper is available from the lead contact upon request.

---

**Figure S1.**
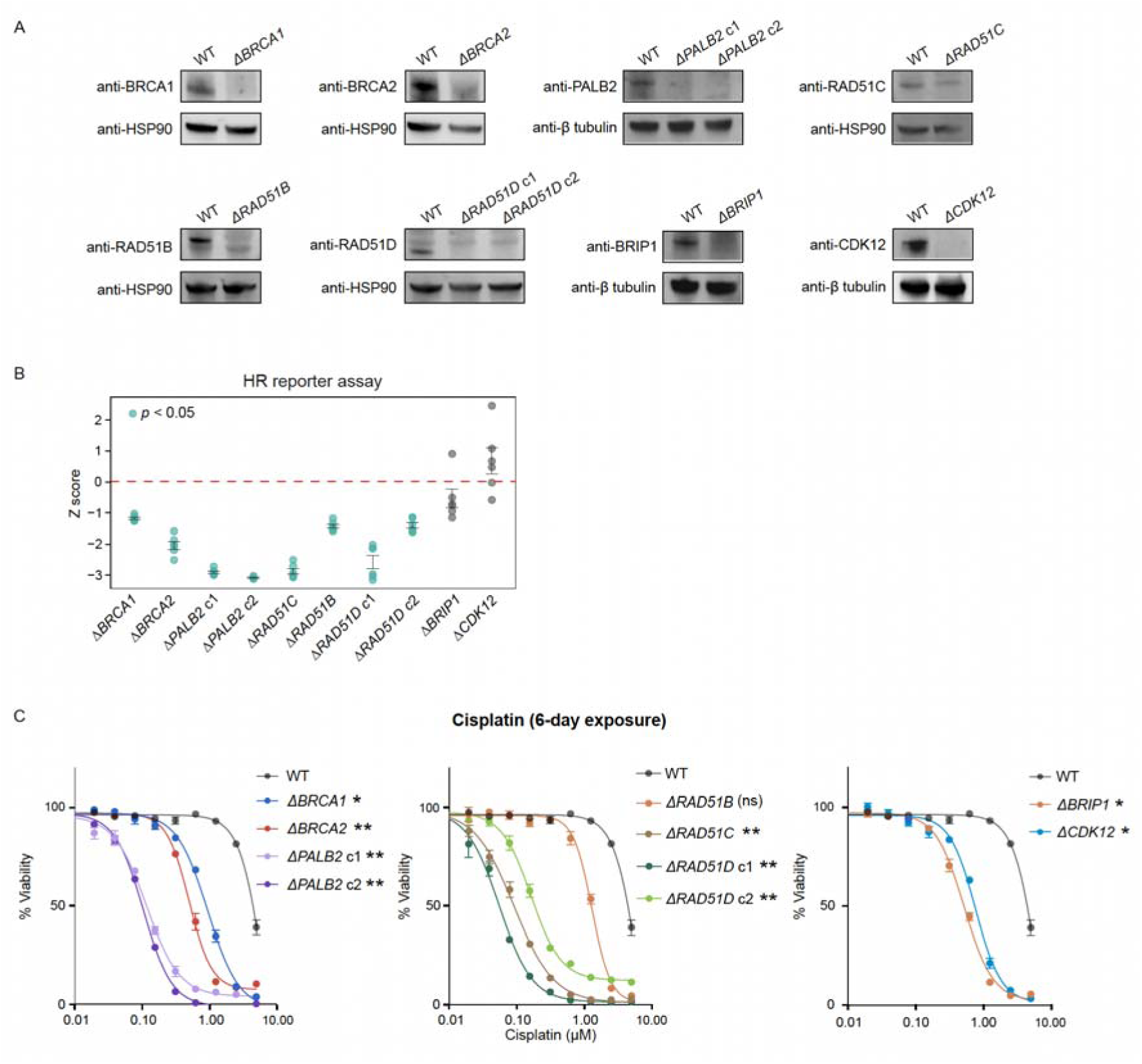
Phenotypic characterizations of isogenic RPE HRd cell lines, related to Figure 1 (A) Immunoblots of isogenic RPE1 HR gene knockouts. (B) HR repair capacity of isogenic HRd gene knockouts as measured using extrachromosomal plasmid reporter assay. Presented as *z* score versus unedited WT. *n*=3 technical replicates. Error bars represent mean_±_SEM. Two-tailed Welch’s t test versus WT; *p*≤0.05 (*), *p*≤0.01 (**). (C) Dose response of isogenic HR gene knockouts to cisplatin (6-day exposure). *n*=3 technical replicates. Error bars represent mean_±_SEM. Two-sided Wilcoxon test versus WT; *p*≤0.05 (*), *p*≤0.01 (**).

**Figure S2.**
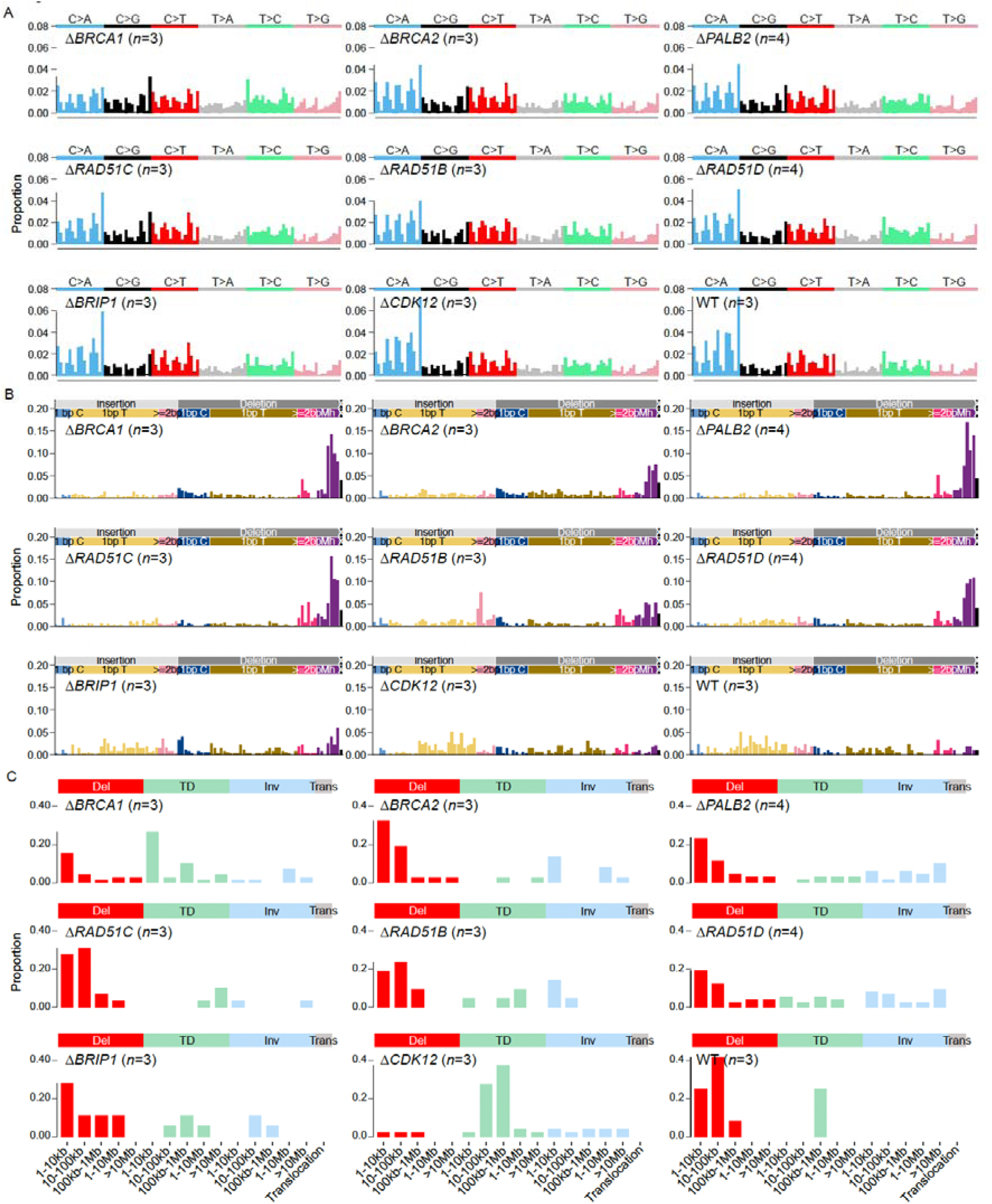
Aggregated genotype-specific SBS, indel, and rearrangement profiles of isogenic RPE1 HRd cell lines, related to Figures 1 and 2 (A) Aggregated genotype-specific SBS profiles of RPE1 HR gene knockouts. (B) Aggregated genotype-specific indel profiles of RPE1 HR gene knockouts. (C) Aggregated genotype-specific rearrangement profiles of RPE1 HR gene knockouts.

**Figure S3.**
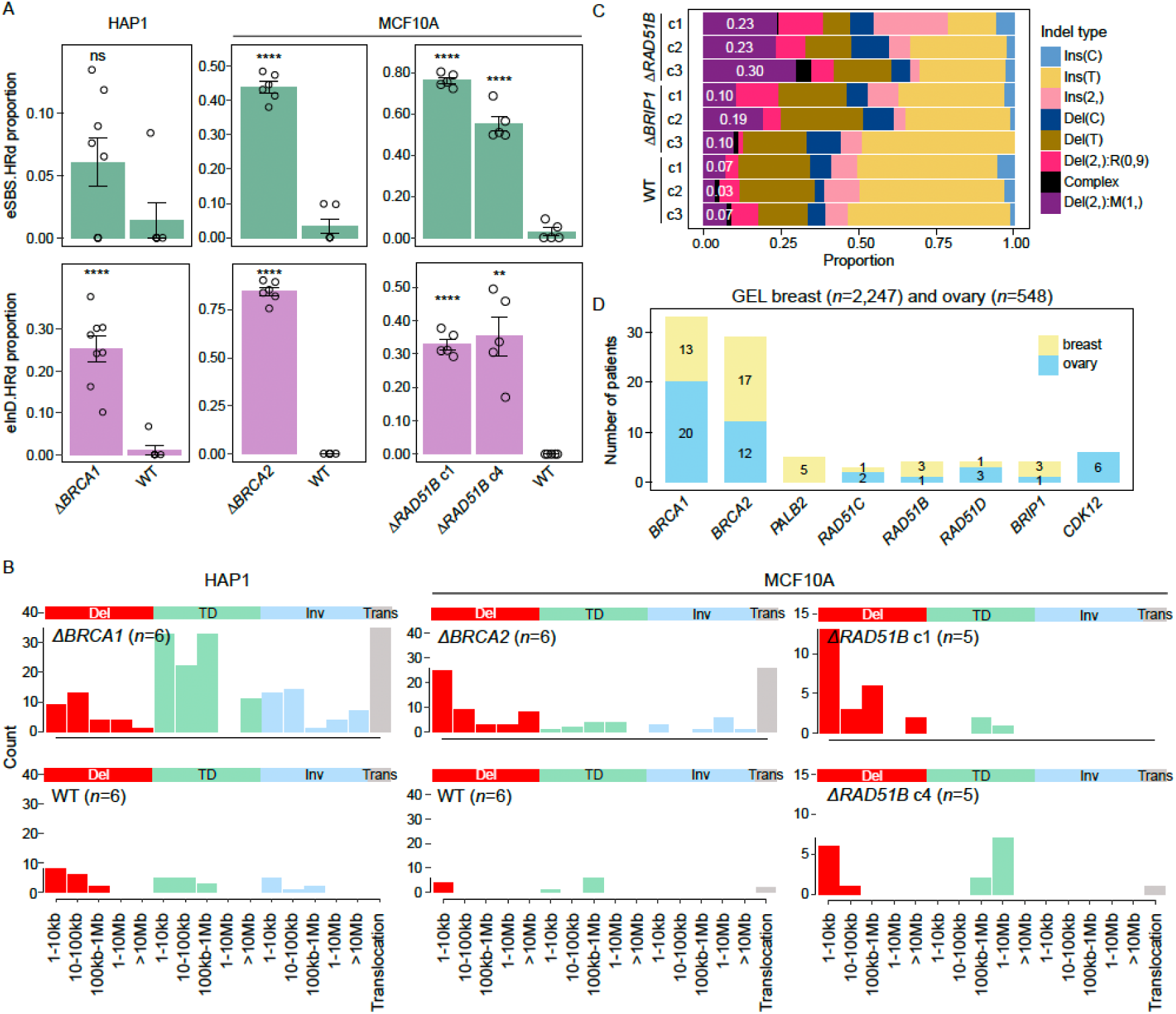
HRd signatures in alternative cellular models and cancers, related to Figure 2 (A) Proportional exposure of RPE1-derived eSBS.HRd and eInD.HRd in HAP1 Δ*BRCA1* (*n*=6), MCF10A Δ*BRCA2* (*n*=6), conditional Δ*RAD51B* (*n*=5 per genotype), and respective unedited WT in each cell line (*n*=3). Error bars represent mean_±_SEM. Two-sided Welch’s t test versus respective WT; *p*≤0.05 (*), *p*≤0.01 (**), *p*≤0.001 (***), *p*≤0.0001 (****). (B) Aggregated rearrangement profiles of HAP1 Δ*BRCA1,* MCF10A Δ*BRCA2*, conditional Δ*RAD51B*, and respective WT per cell line. *n* denotes the number of subclones per genotype. (C) Relative proportions of indel mutation subtypes per subclone of RPE1 *ΔRAD51B, ΔBRIP1,* and WT. Numbers indicate the proportional microhomology-mediated deletions. (D) Prevalence of biallelic *loss-of-function* HR gene mutations (*n*=8 genes in this manuscript) in GEL breast (*n*=2,247) and ovarian (*n*=548) cohorts.

**Figure S4.**
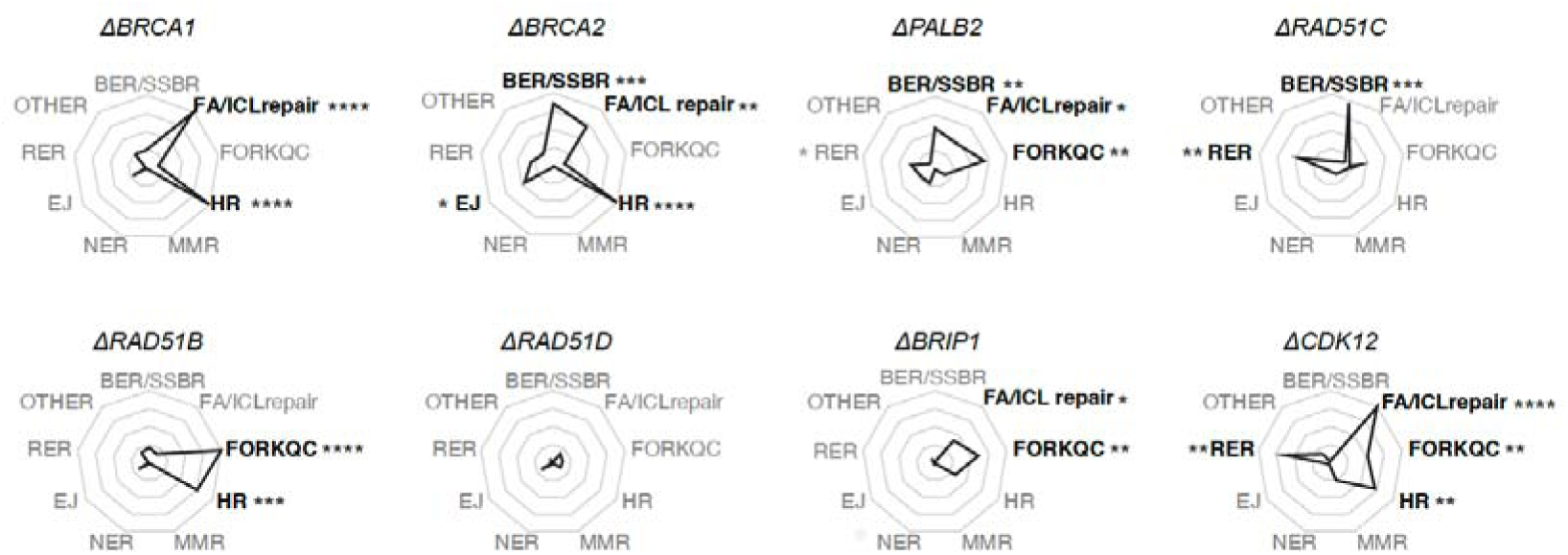
DNA repair pathway enrichment among genotype-specific synthetic lethal targets in RPE1 isogenic knockouts, related to Figure 3. Bold texts indicate the most significantly enriched pathways. Fisher’s exact test; *p*≤0.05 (*), *p*≤0.01 (**), *p*≤0.001 (***), *p*≤0.0001 (****).

**Figure S5.**
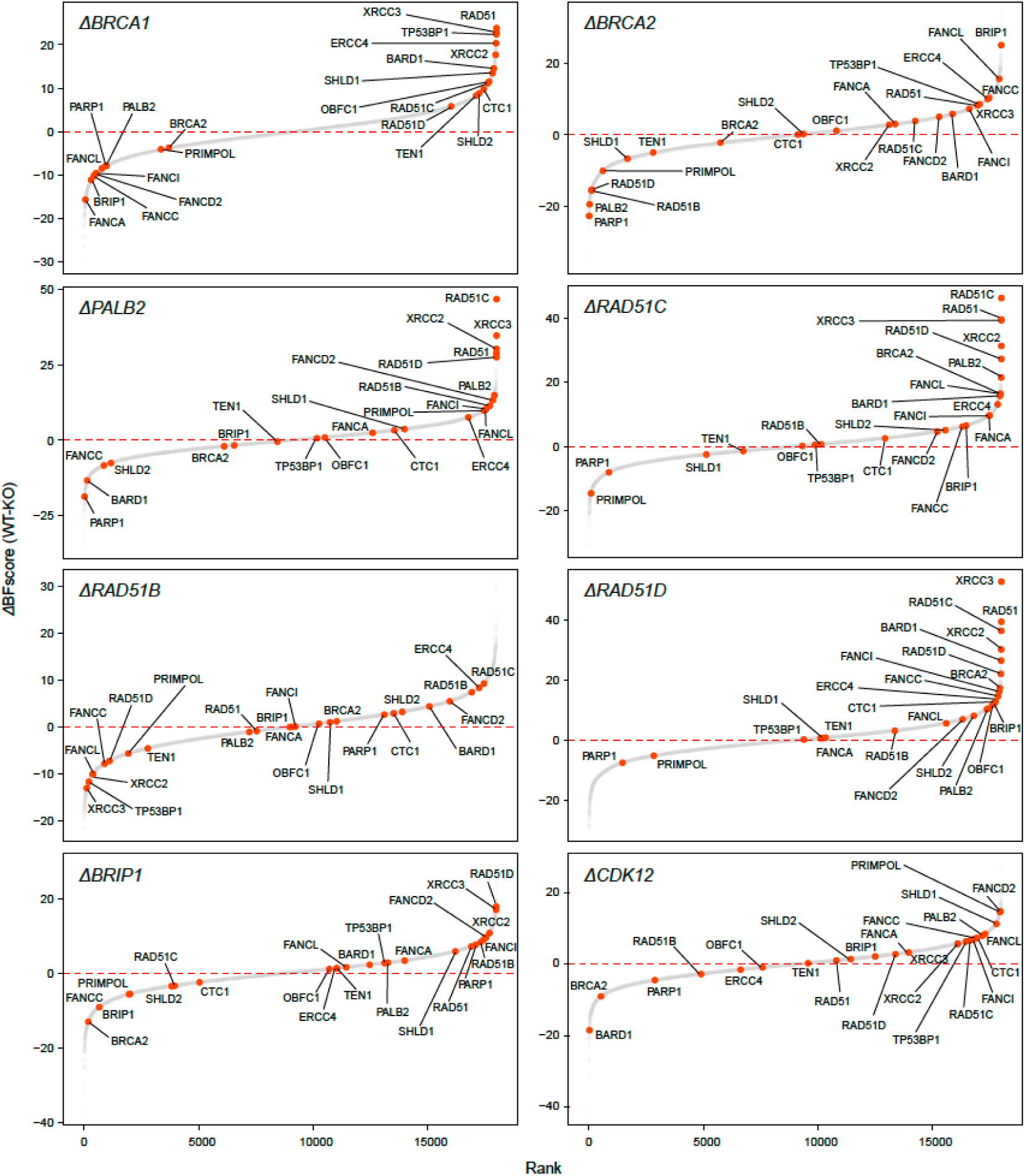
ΔBF scores rank plots of RPE1 isogenic knockout CRISPR screens, related to Figure 3. Delta Bayes factor (ΔBF) scores represent the difference in viability between WT and knockout cells (WT-knockout).

**Figure S6.**
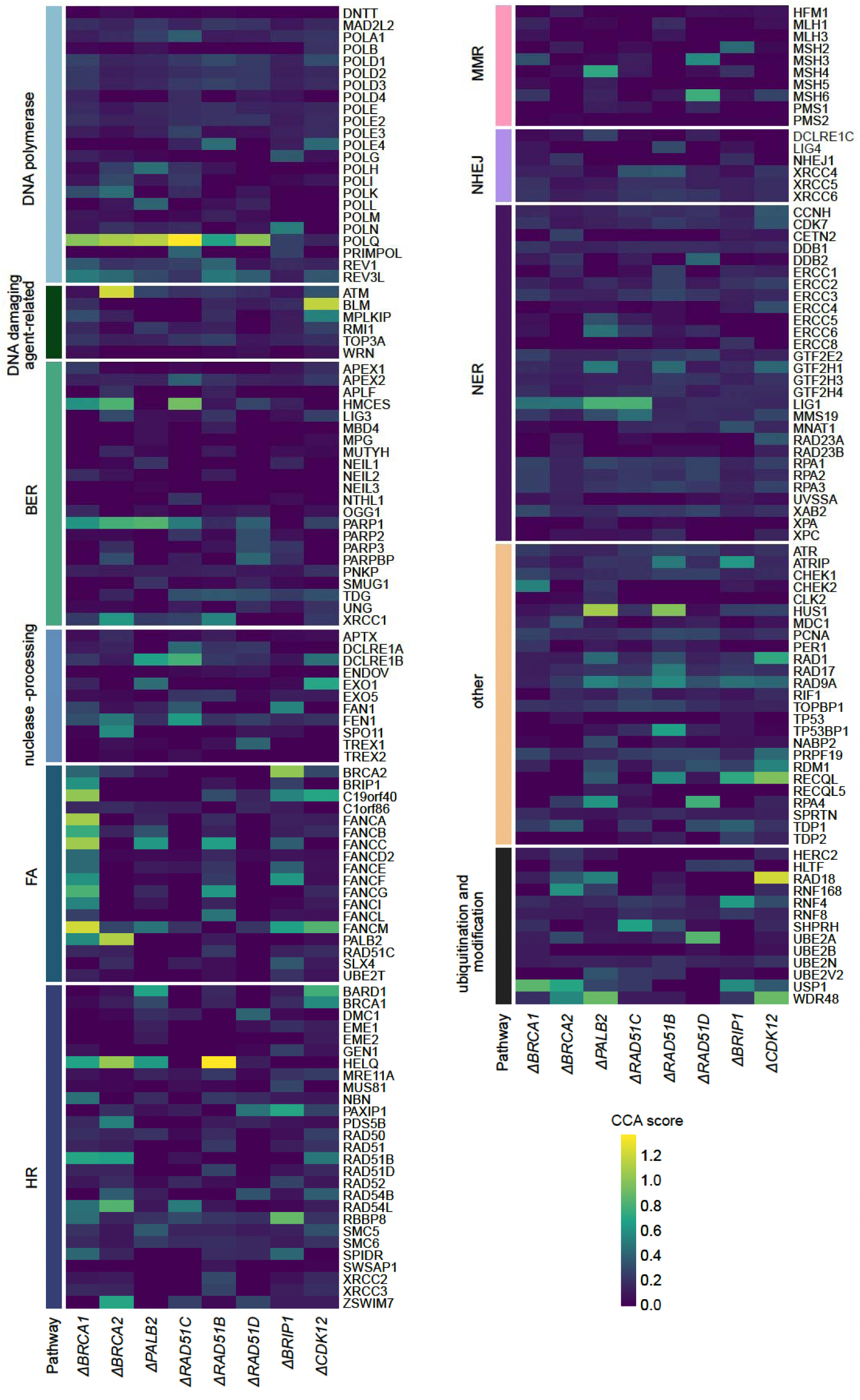
Heatmap of CCA scores of RPE1 isogenic knockout CRISPR screens, related to Figure 3. Genes are grouped by DNA repair pathways. Higher CCA scores indicate stronger genotype-selective synthetic lethal effects.

**Figure S7.**
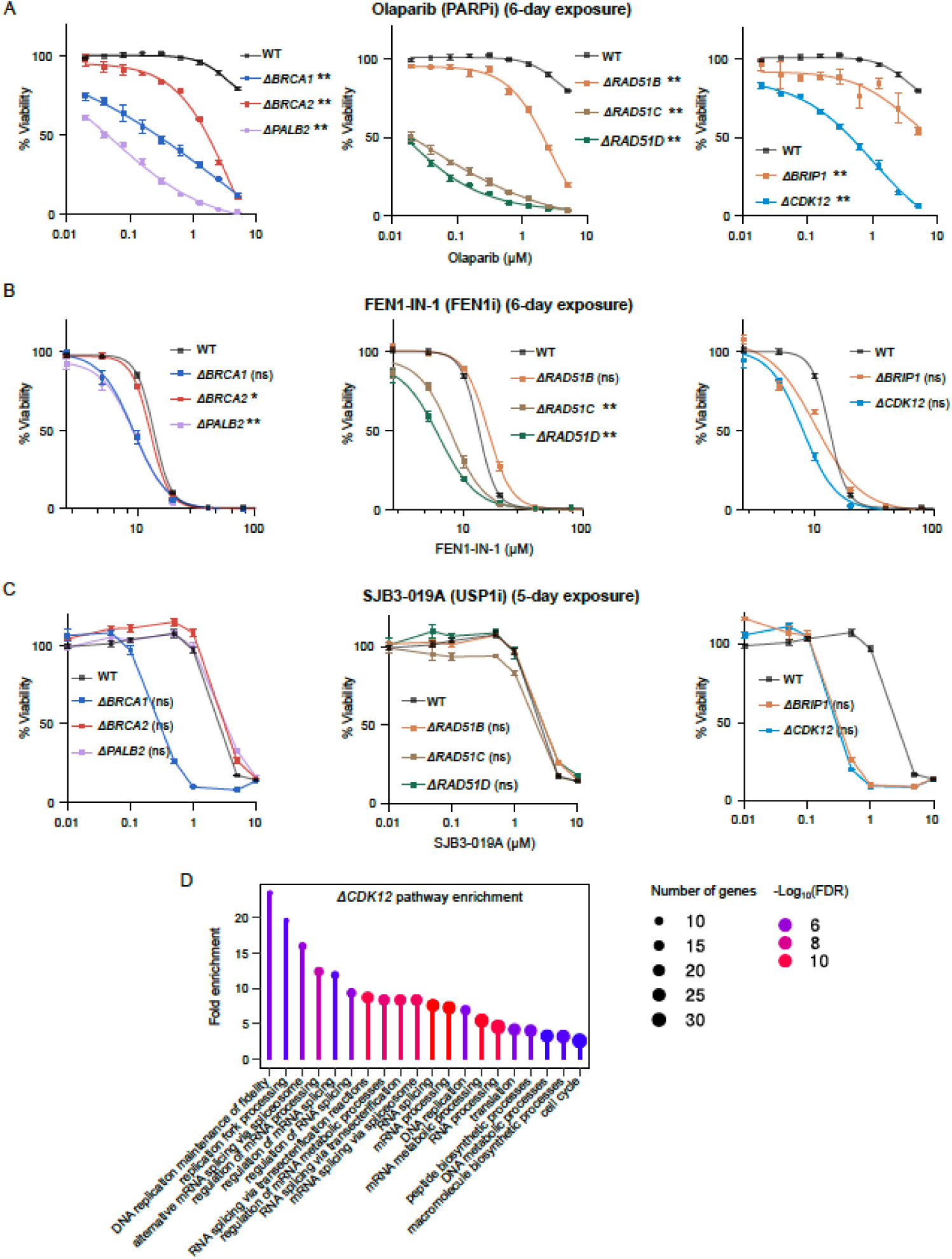
Validation of selected HRd SL targets, related to Figure 3 (A) Dose response of RPE1 isogenic knockouts to PARP inhibitor, olaparib. *n*=3 technical replicates. Error bars represent mean_±_SEM. Two-sided Wilcoxon test versus WT; *p*≤0.05 (*), *p*≤0.01 (**). (B) Dose response of RPE1 isogenic knockouts to FEN1 inhibitor, FEN1-IN-1. *n*=3 technical replicates. Error bars represent mean]±]SEM. Two-sided Wilcoxon test versus WT; *p*≤0.05 (*), *p*≤0.01 (**). (C) Dose response of RPE1 isogenic knockouts to USP1 inhibitor, SJB3-019A. *n*=3 technical replicates. Error bars represent mean]±]SEM. Two-sided Wilcoxon test versus WT. (D) Top upregulated biological pathways among *n*=2,009 differentially expressed genes (Benjamin-Hochberg adjusted *p*<0.05 and L_2_FC≥0.6) comparing RPE1 Δ*CDK12* cells (*n*=3) to WT cells (*n*=3).

**Figure S8.**
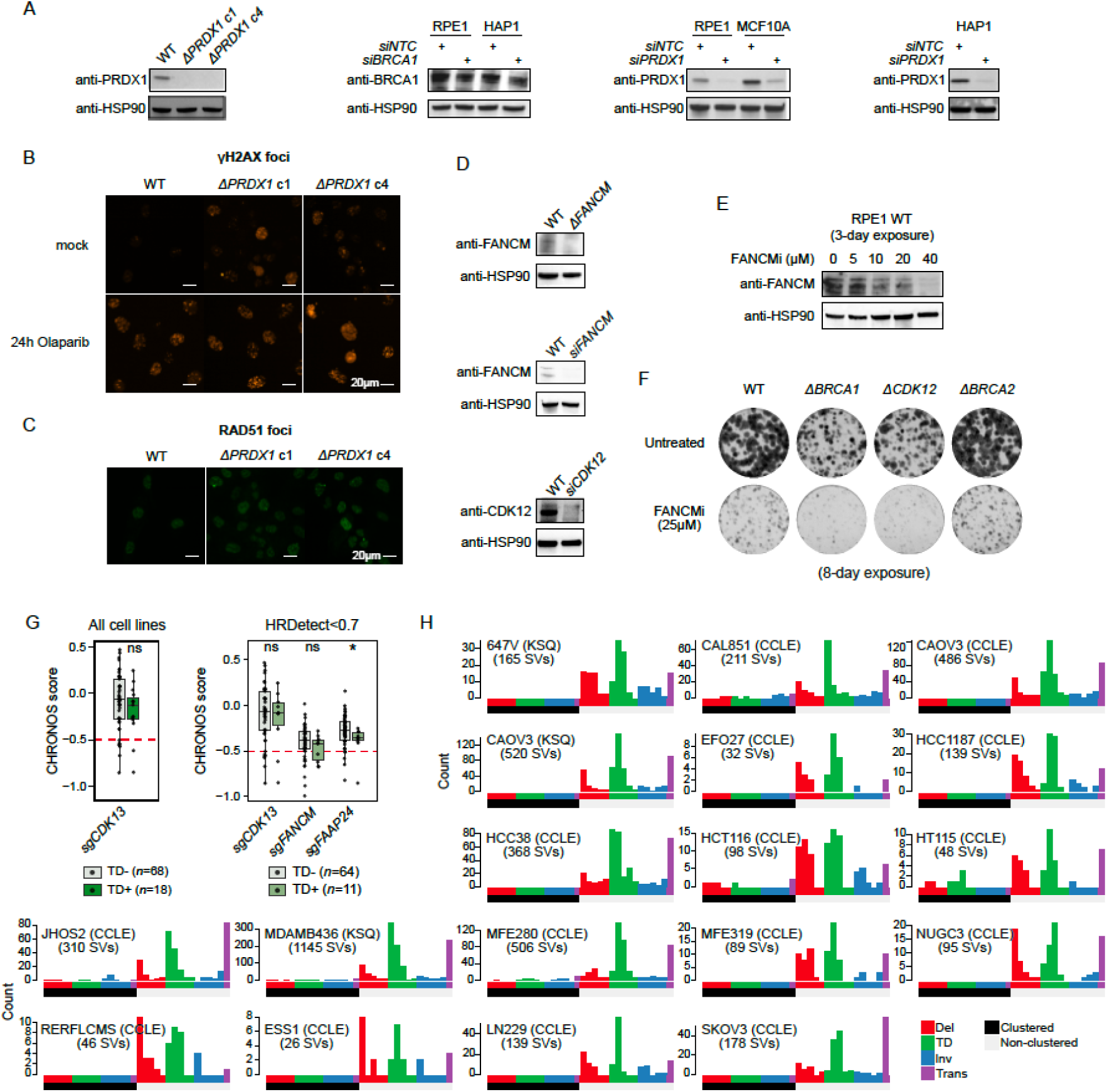
Validating *PRDX1*: HRd and *FANCM:CDK12* synthetic lethal interactions, related to Figures 4 and 5 (A) Immunoblots of RPE1 Δ*PRDX1* (clones c1 and c4), siRNA knockdown (3 days post-transfection) of *PRDX1* in RPE1, MCF10A, and HAP1 cells, related to Figure 4B. (B) Representative immunofluorescence images of γH2AX foci in RPE1 WT and Δ*PRDX1* ± 24h 5μM olaparib exposure, related to Figure 4E. γH2AX measures DNA double strand breaks. (C) Representative immunofluorescence images of baseline RAD51 foci in RPE1 WT and Δ*PRDX1*. Related to Figure 4H. (D) Immunoblots of RPE1 Δ*FANCM* and siRNA knockdown of *FANCM* and *CDK12*, related to Figure 5F. (E) Immunoblot of *FANCM* in RPE1 WT cells treated with different concentrations of FANCM inhibitor, FANCM-BTR PPI-IN-1 (3-day exposure), related to Figure 5G. (F) Representative images of colony formation assay of RPE1 WT, Δ*BRCA1,* Δ*CDK12,* and Δ*BRCA2* following 8-day treatment with FANCM inhibitor, FANCM-BTR PPI-IN-1 (25μM), related to Figure 5G. (G) CHRONOS essentiality scores of *CDK13*, *FANCM,* and *FAAP24* derived from the BROAD DepMap project. Cell lines were fitted with rearrangement signatures and grouped according to whether they harbored TD signatures (R1, R3, R14) (TD+) or not (TD-). Left: all cell lines that satisfy the above criteria. Right: cell lines with HRDetect probability ≥0.7 were excluded. Two-sided Welch’s t-tests, TD+ versus TD-; *p*≤0.05 (*). Related to Figure 5H. (H) Rearrangement profiles of whole-genome sequenced DepMap *n*=18 TD+ cell lines. WGS data source for each cell line is indicated in parentheses (CCLE, Cancer Cell Line Encyclopedia; KSQ, KSQ Therapeutics, Inc.). Related to Figure 5H.

**Figure S9.**
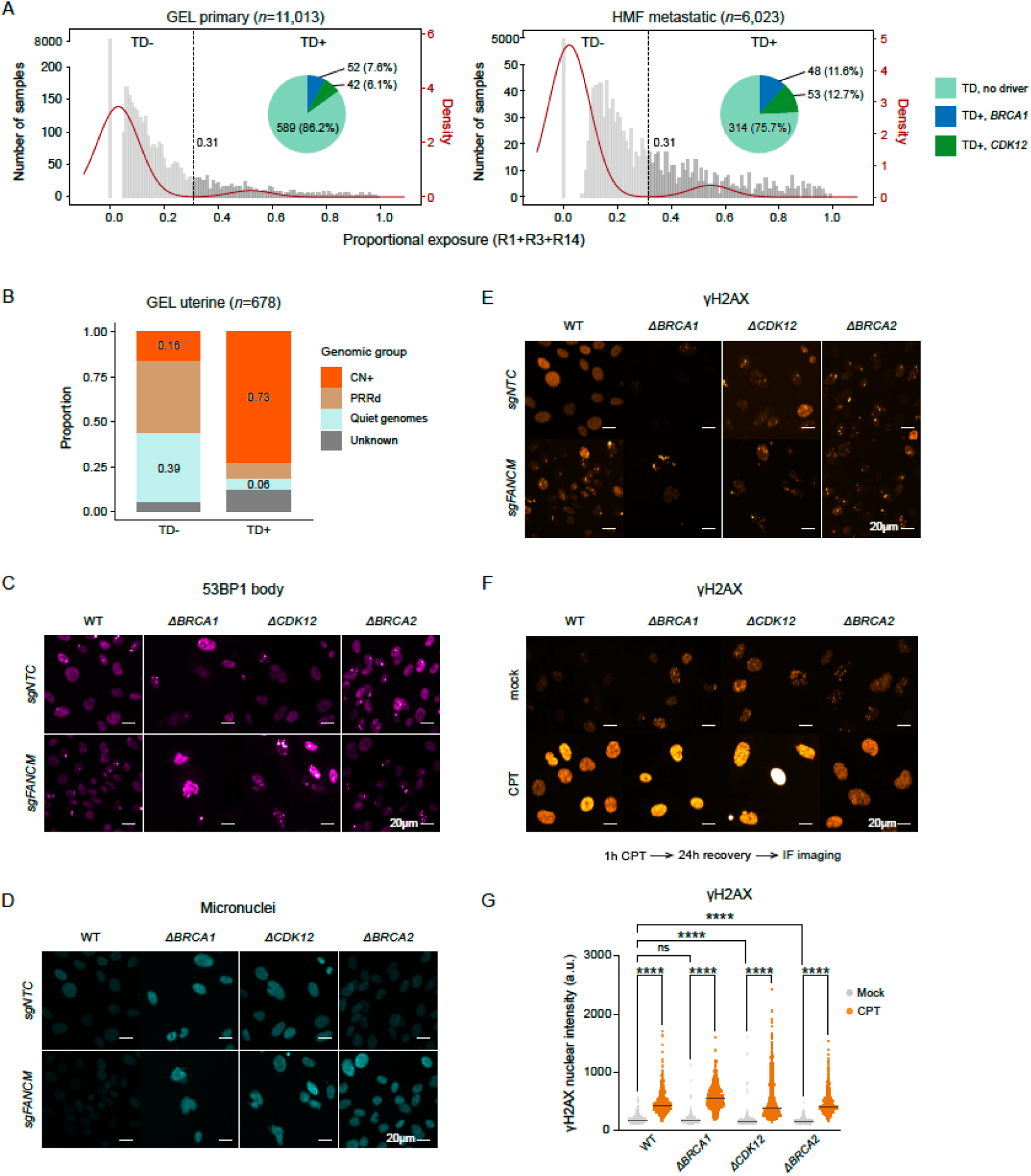
TD phenotype and TD:*FANCM* SL, related to Figures 5 and 6 (A) Histograms of proportional exposures of TD mutational signatures (R1+R3+R14) in *n*=11,013 GEL primary and *n*=6,023 HMF metastatic pan-cancer cohorts. Red curves indicate fitted Gaussian mixture models in each cohort. Vertical dashed lines mark thresholds distinguishing TD+ versus TD-samples. Pie charts demonstrate TD+ samples with and without *BRCA1* or *CDK12* biallelic *loss-of-function* mutations. Related to Figure 5I. (B) Distribution of copy-number phenotypes among TD− and TD+ tumours in the GEL uterine cancer cohort (*n* = 678). Bars show the proportion of tumors in each TD group classified as copy-number-high (CN+), post-replicative repair deficient (PRRd), copy-number quiet, or unknown copy number status. TD status was assigned using the threshold defined in (A). Samples with unknown copy-number status were retained in this analysis but excluded from the survival analysis in Figure 5J. (C) Representative immunofluorescence images of 53BP1 body in RPE1 WT, Δ*BRCA1*, Δ*CDK12* and Δ*BRCA2* with and without *FANCM* depletion. Related to Figure 6A. (D) As in (C), for micronuclei. Related to Figure 6B. (E) As in (C), for γH2AX. Related to Figure 6C. (F) Representative immunofluorescence images of γH2AX nuclear intensity in RPE1 WT, Δ*BRCA1*, Δ*CDK12,* and Δ*BRCA2* 24h following 1h exposure to 1uM CPT and 24h recovery. (G) Quantification of γH2AX nuclear intensity in (F). Center lines indicate the median value. Two-way ANOVA; *p*≤0.05 (*), *p*≤0.01 (**), *p*≤0.001 (***), *p*≤0.0001 (****).

**Figure S10.**
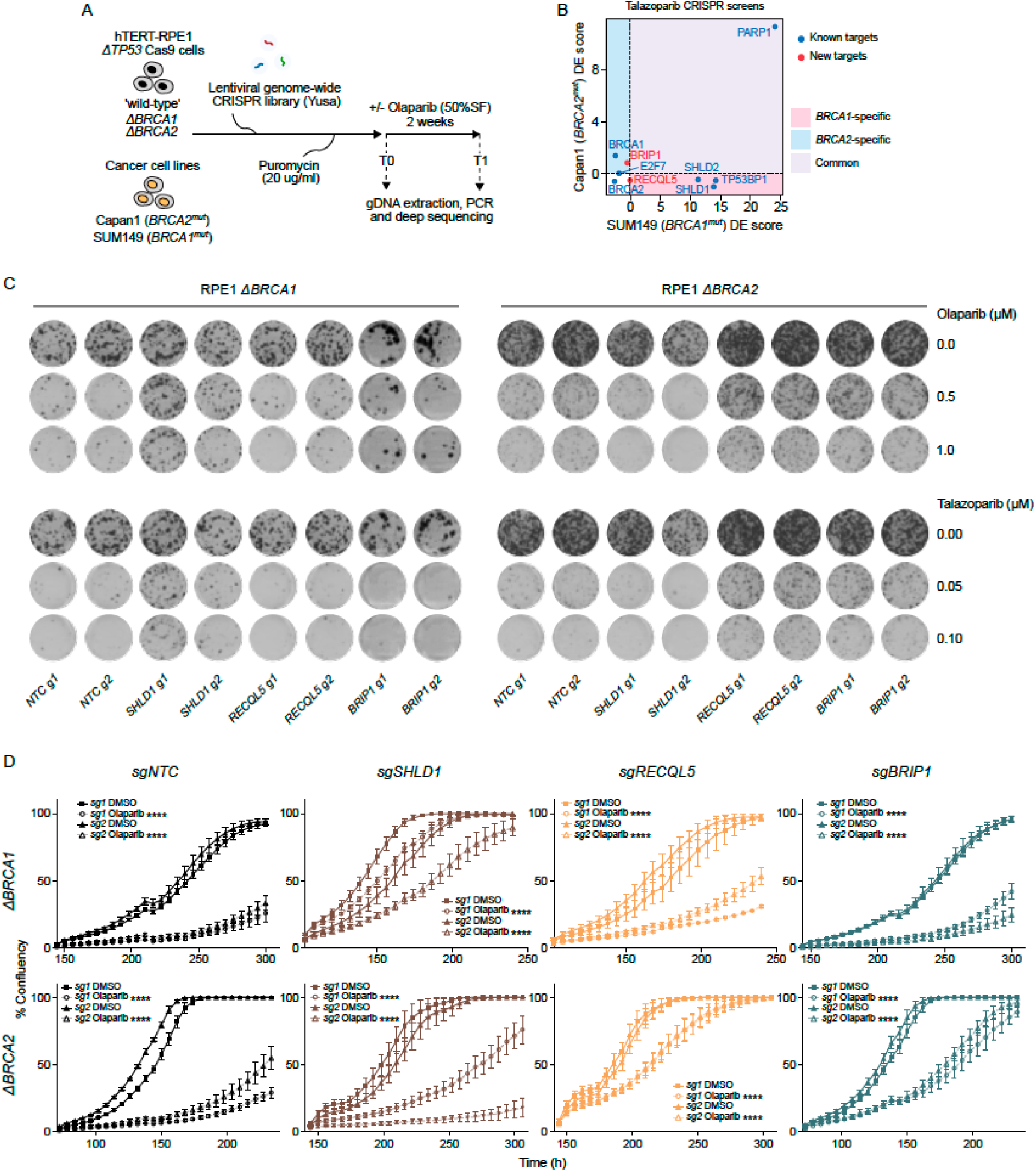
PARPi resistance CRISPR screens identify genotype-specific resistance drivers, related to Figure 7 (A) Schematic of genome-wide CRISPR dropout screens ± PARPi selection in isogenic RPE1 knockouts (*i.e.,* Δ*TP53* (WT), Δ*BRCA1*, Δ*BRCA2*) (olaparib) and HRd cancer cell lines (*i.e., BRCA1^mut^* SUM149; *BRCA2^mut^* Capan1) (olaparib or talazoparib). (B) DE scores of known PARPi resistance genes and new *BRCA2*-specific PARPi resistance genes in talazoparib CRISPR resistance screens of *BRCA1^mut^* SUM149 and *BRCA2^mut^* Capan1. (C) Representative images of colony formation of RPE1 Δ*BRCA1* and Δ*BRCA2* in PARPi (olaparib or talazoparib) for 14 days, with and without *sgRNA* knockout of *SHLD1*, *RECQL5*, and *BRIP1*. (D) Proliferation of RPE1 Δ*BRCA1* and Δ*BRCA2* in olaparib or DMSO, with and without *sgRNA* knockout of *SHLD1*, *RECQL5*, and *BRIP1*. *n*=4 technical replicates. Error bars represent mean]±]SEM Statistical comparisons were made between olaparib-treated and respective DMSO-treated conditions. Two-way ANOVA; *p*≤0.05 (*), *p*≤0.01 (**), *p*≤0.001 (***), *p*≤0.0001 (****).

